# Comparative multiplexed interactomics of SARS-CoV-2 and homologous coronavirus non-structural proteins identifies unique and shared host-cell dependencies

**DOI:** 10.1101/2020.07.13.201517

**Authors:** Jonathan P. Davies, Katherine M. Almasy, Eli F. McDonald, Lars Plate

## Abstract

Human coronaviruses (hCoV) have become a threat to global health and society, as evident from the SARS outbreak in 2002 caused by SARS-CoV-1 and the most recent COVID-19 pandemic caused by SARS-CoV-2. Despite high sequence similarity between SARS-CoV-1 and −2, each strain has distinctive virulence. A better understanding of the basic molecular mechanisms mediating changes in virulence is needed. Here, we profile the virus-host protein-protein interactions of two hCoV non-structural proteins (nsps) that are critical for virus replication. We use tandem mass tag-multiplexed quantitative proteomics to sensitively compare and contrast the interactomes of nsp2 and nsp4 from three betacoronavirus strains: SARS-CoV-1, SARS-CoV-2, and hCoV-OC43 – an endemic strain associated with the common cold. This approach enables the identification of both unique and shared host cell protein binding partners and the ability to further compare the enrichment of common interactions across homologs from related strains. We identify common nsp2 interactors involved in endoplasmic reticulum (ER) Ca^2+^ signaling and mitochondria biogenesis. We also identifiy nsp4 interactors unique to each strain, such as E3 ubiquitin ligase complexes for SARS-CoV-1 and ER homeostasis factors for SARS-CoV-2. Common nsp4 interactors include *N*-linked glycosylation machinery, unfolded protein response (UPR) associated proteins, and anti-viral innate immune signaling factors. Both nsp2 and nsp4 interactors are strongly enriched in proteins localized at mitochondrial-associated ER membranes suggesting a new functional role for modulating host processes, such as calcium homeostasis, at these organelle contact sites. Our results shed light on the role these hCoV proteins play in the infection cycle, as well as host factors that may mediate the divergent pathogenesis of OC43 from SARS strains. Our mass spectrometry workflow enables rapid and robust comparisons of multiple bait proteins, which can be applied to additional viral proteins. Furthermore, the identified common interactions may present new targets for exploration by host-directed anti-viral therapeutics.

## INTRODUCTION

Coronaviruses (CoVs) are positive single-stranded RNA viruses capable of causing human disease with a range of severity. While some strains, such as endemic hCoV-OC43, cause milder common-cold like symptoms, other strains are associated with more severe pathogenesis and higher lethality, including SARS-CoV-1 (emerged in 2002), MERS-CoV (in 2012), and most recently SARS-CoV-2, the causative agent of COVID-19 ^1,2^. Despite the relevance of CoVs for human health, our understanding of the factors governing their divergent pathogenicity remains incomplete. Pathogenicity may be mediated by a variety of factors, including different specificities and affinity for different cell surface receptors such as angiotensin-converting enzyme 2 (ACE2) for SARS-CoV-1 and SARS-CoV-2 ^1,3^ or 9-*O*-acetylated sialic acid for hCoV-OC43^4^. CoV strains also engage a variety of host immune processes in infected cells. Pathogenic strains more strongly interfere with interferon I signaling ^4,5^ and induce apoptosis and pyroptosis ^6–9^. Ensembles of virus-host protein-protein interactions (PPIs) orchestrate the reprogramming of these processes during infection.

Coronaviruses possess the largest known RNA viral genomes, approximately 30kbp in length. The 5’ 20 kb region of the genome encodes for two open reading frames (orf1a/1ab) that produce 16 non-structural proteins (nsp1-nsp16) needed to form the viral replication complex, while the 3’ proximal region encodes for the structural proteins and several accessory factors with varying roles (Figure 1A). Previous protein-protein interaction studies of individual CoV proteins have shed light on their functions in the infected host cells and putative roles during pathogenesis. Yeast-two hybrid studies of coronavirus proteins have identified intraviral interactions^10^ and interactions between nsp1 and immunophilins^11^, and a proximity-labeling approach was used to determine host proteins concentrated in sites of replication^12^.

**Figure 1.**
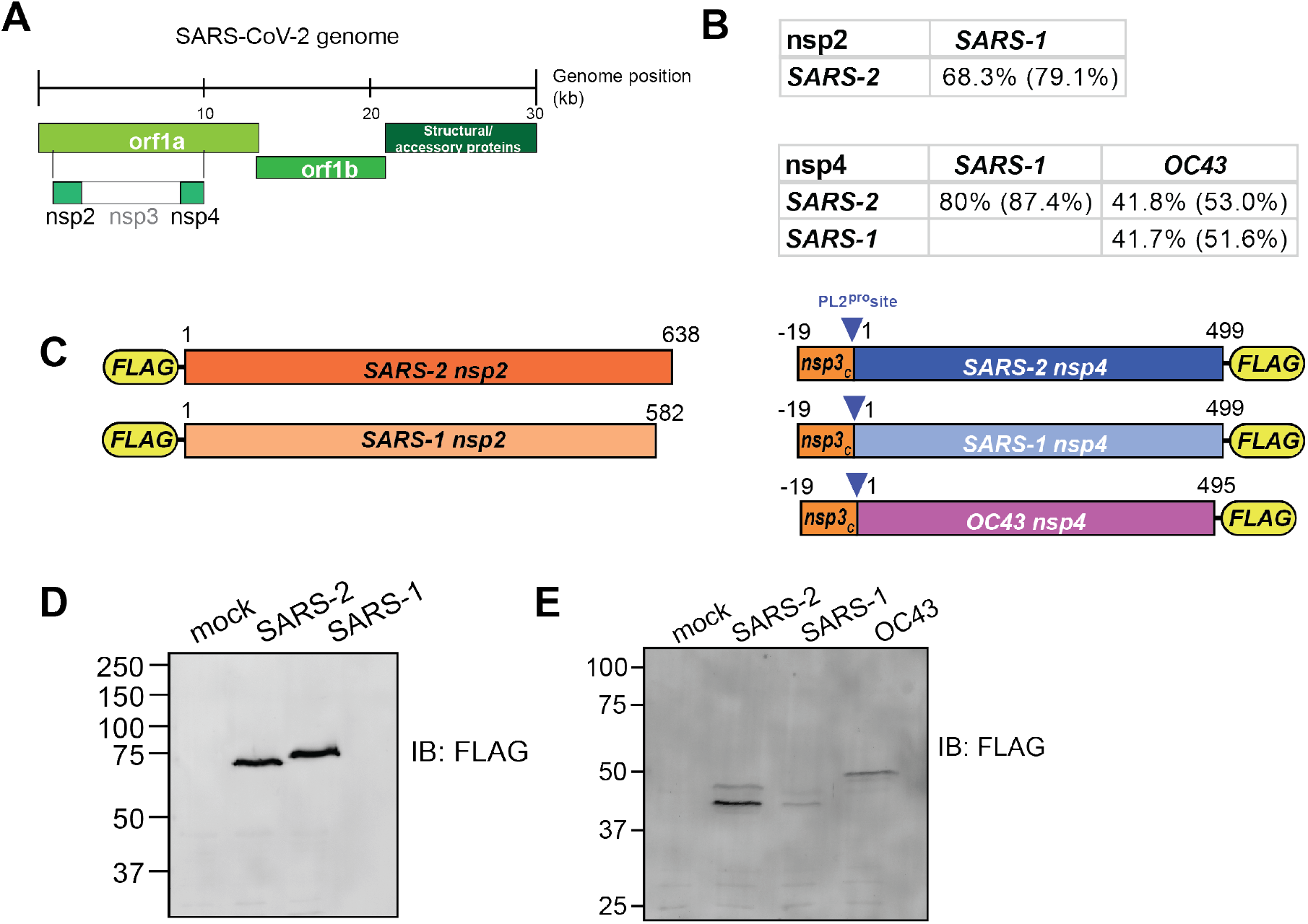
Design and validation of expression of CoV nsp2 and nsp4 constructs for affinity purification. (A) Schematic of SARS-CoV-2 genome organization. (B) Amino acid sequence identity and similarity (in parentheses) for comparisons of nsp2 and nsp4 homologs. Sequence alignments are shown in Figure S1A-B. (C) Nsp2 and nsp4 FLAG-tagged construct designs. Nsp2 constructs contain an N-terminal FLAG-tag. Nsp4 constructs contain a 19 amino acid leader sequence from nsp3 at the N-terminus, including the PL2^pro^ cleavage site, along with a C-terminal FLAG-tag. (D-E) Western blot of nsp2 and nsp4 homologs expressed in HEK293T cells. Cell were transiently transfected with FLAG-nsp2 (D) or nsp4-FLAG (E). Proteins were detected using an anti-FLAG antibody.

Affinity purification-mass spectrometry (AP-MS) is a powerful tool to study virus-host interactions and has been used extensively to examine how viruses reorganize host cells^13–16^. A prior AP-MS study of SARS-CoV-1 nsp2 identified multiple host interactors including prohibitin 1/2 (PHB1/2)^17^. Most notably, Gordon et al. recently profiled host interactors for 26 SARS-CoV-2 proteins^18^. While these studies enabled important insight on individual viral protein functions, they focused on single CoV strains, limiting direct cross-strain comparisons.

Here, we sought to profile and compare the host interaction profiles of nsps from multiple hCoVs, namely hCoV-OC43, SARS-CoV-1, and SARS-CoV-2. Through comparative interactomics, we identify both conserved and unique interactors across various strains. Notably, quantitative analysis of interaction enrichment enables nuanced differentiation between shared interactions for each coronavirus protein. Through this approach we discovered both conserved and novel functions of viral proteins and the pathways by which they manipulate cellular processes. Comparisons across strains may also provide clues into the evolutionary arms race between virus and host proteins to hijack or protect protein-protein interfaces^19^. Additionally, identified host dependencies can potentially be exploited as targets for host-directed antiviral therapeutics.

In particular, we focus on the host interactors of nsp2 and nsp4. Nsp2 has been suggested to play a role in modifying the host cell environment, although its precise function remains unknown^17^. Nsp2 is dispensable for infection in SARS-CoV-1 ^20^ and has pronounced amino acid sequence differences across coronavirus strains (Figures 1B, S1A). Additionally, early sequence analysis of SARS-CoV-2 identified regions of positive selection pressure in nsp2^21^. Given the variability of sequence across strains and the ambiguous function, a comparison of interaction profiles across strains can yield insights into the role of nsp2. In contrast, the role of nsp4, a transmembrane glycoprotein, is better defined, most notably in formation of the double-membrane vesicles associated with replication complexes^22,23^. Unlike nsp2, nsp4 has a high degree of sequence similarity across human coronavirus strains (Figures 1B, S1B).

In this study, we use affinity purification-proteomics to identify interactors of nsp2 from two human coronaviruses (SARS-CoV-1 and SARS-CoV-2) and interactors of nsp4 from three strains (OC43, SARS-CoV-1, and SARS-CoV-2). Quantitative comparative analysis of nsp2 interactors identifies common protein binding partners, including the ERLIN1/2 complex and prohibitin complex involved in regulation of mitochondrial function and calcium flux at ER-mitochondrial contact sites. We also identify overlapping nsp4 interactors, including *N*-linked glycosylation machinery, UPR associated factors, and anti-viral innate immune signaling proteins. Unique interactors of different nsp4 homologs include E3 ubiquitin ligase complexes for SARS-CoV-1, and ER homeostasis factors for SARS-CoV-2. In particular, we found nsp2 and nsp4 interactors are strongly enriched for mitochondrial-associated ER membranes (MAM) factors, suggesting a potential mechanism to affect calcium homeostasis and other host processes at these organelle contact sites.

## RESULTS

### Design and validation of expression of CoV nsp2 and nsp4 constructs for affinity purification

The two main open reading frames of the CoV viral genome, orf1a and orf1ab, encode for 16 non-structural proteins which perform a variety of tasks during the infection cycle (Figure 1A). We focus our analysis on two of these proteins, nsp2 and nsp4. Nsp2 is a less functionally-well understood protein with less than 70% amino acid sequence identity between the SARS-CoV-1 and SARS-CoV-2 homologs (Figures 1B, S1A). Nsp4 is a component of the CoV replication complex that is 80 % identical between SARS strains, but only 42% between SARS and OC43 strains, a less clinically severe human CoV (Figures 1B, S1B).

To compare the virus-host protein-protein interactions of nsp2 and nsp4 across multiple CoV strains, we designed FLAG-tagged expression constructs for affinity purification (Figure 1C). SARS-CoV-1 and SARS-CoV-2 nsp2 constructs contain an N-terminal FLAG-tag, while the SARS-CoV-1, SARS-CoV-2, and OC43 nsp4 constructs contain a C-terminal FLAG-tag. In addition, nsp4 constructs contain a 19 amino acid leader sequence corresponding to the C-terminus of nsp3, which includes the nsp3-PL2^pro^ cleavage site for proper nsp4 translocation into the ER membrane^24,25^.

Protein constructs were transiently transfected into HEK293T cells and proteins were detected by immunoblotting for the FLAG-tag. While HEK293T cells are not representative of the primary physiological target tissue, these cells are permissive to infection, and were able to recapitulate strong interactors expected in lung tissue in a prior SARS-CoV-2 interactome study^18^. The nsp2 constructs were detectable as a single protein band at the expected molecular weight (Figure 1D), while nsp4 constructs displayed two distinct bands at a lower size than its expected molecular weight (Figure 1E). This lower apparent molecular weight was previously reported and the different bands likely correspond to different glycosylation states^18^. To ensure nsp4 is expressed fully and the detected products do not correspond to a truncated protein, we immunopurified the protein using FLAG-agarose beads and analyzed the purified protein by LC-MS. We detected peptide fragments spanning the N- and C-termini with overall sequence coverage of up to 62% (Figure S1C-E) confirming expression of the full proteins.

### Affinity purification-mass spectrometry identifies nsp2 interactors

To identify host cell interaction partners of the distinct CoV non-structural proteins, we employed an affinity purification-mass spectrometry workflow (Figure 2A). The protein constructs were expressed in HEK293T cells, gently lysed in mild detergent buffer, and co-immunopurified from whole cell lysates using anti-FLAG agarose beads. Virus-host protein complexes were then reduced, alkylated, and trypsin digested. Importantly, we used tandem mass tag (TMT)-based multiplexing using TMTpro-16plex or TMT-11plex for relative quantification of protein abundances. For this purpose, 4 – 6 co-immunoprecipitation replicates for respective nsp2 homologs were pooled into a single MS run. Co-immunoprecipitations from mock GFP transfected cells were included to differentiate non-specific background proteins (Figure 2B). Overall, the data set included three individual MS runs containing 34 Co-IP (SARS-CoV-2 n = 13; SARS-CoV-1 n = 9; GFP (mock) n = 12) (Figure S2A).

**Figure 2.**
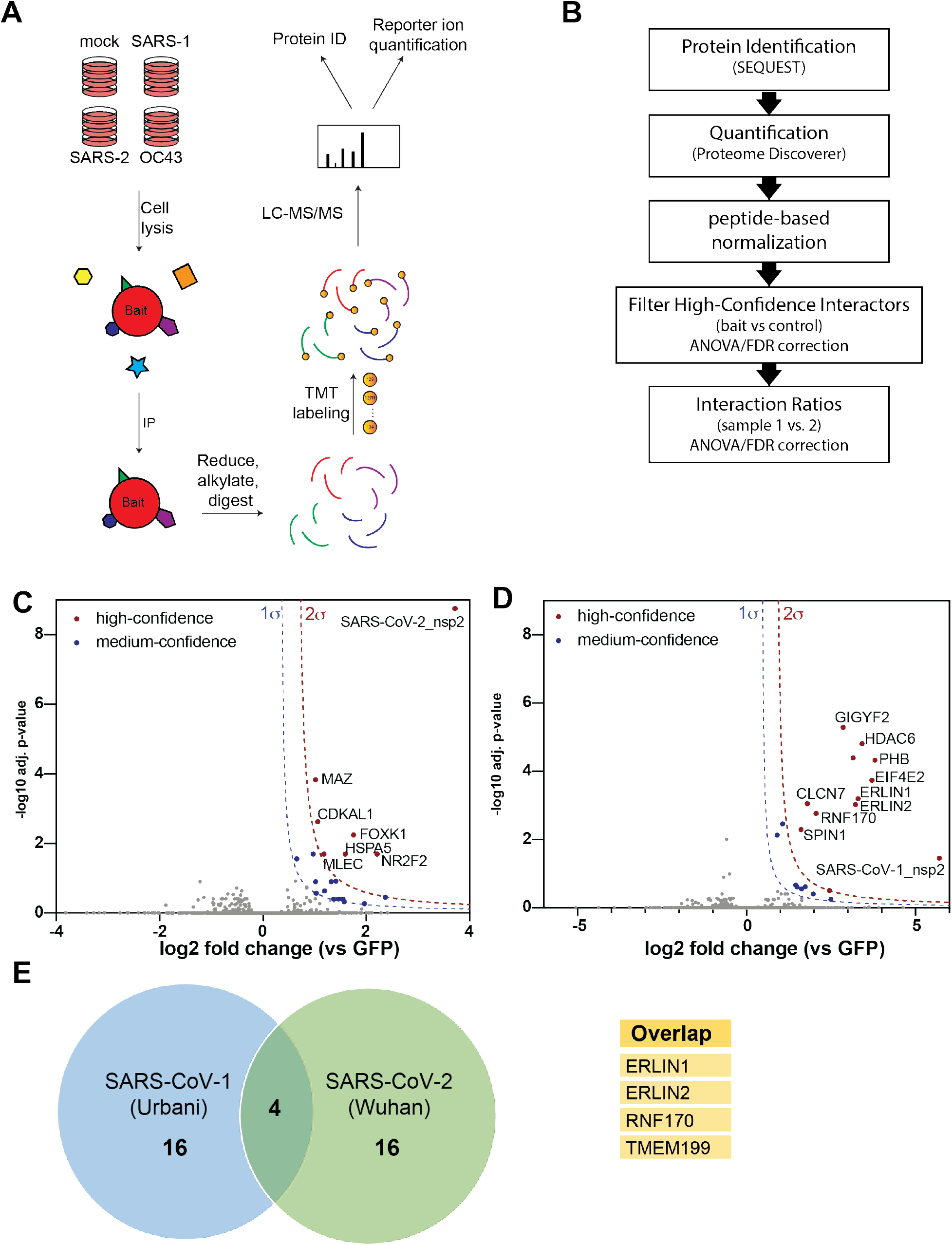
Affinity purification-mass spectrometry (AP-MS) identifies nsp2 interactors. (A) General AP-MS workflow to quantitatively determine interactors of viral nsp homolog. HEK293T cells are transfected with FLAG-tagged expression constructs of nsps as bait or GFP (mock) and lysed. Bait proteins are immunoprecipitated (IP) along with interacting proteins, reduced, alkylated, and tryptic digested. Peptides are then tandem-mass tag (TMT) labeled, pooled, and analyzed by LC-MS/MS for identification and quantification. (B) Data processing workflow. Peptide spectra are identified and matched to corresponding proteins (SEQUEST HT), then quantified based on TMT reporter ion intensity (Proteome Discoverer 2.4). High confidence interactors are filtered by comparing bait vs control. Interaction ratios between bait homologs are determined (log2 fold change) and adjusted p-value calculated using ANOVA. (C-D) Volcano plot of SARS-CoV-2 nsp2 (C) and SARS-CoV-1 nsp2 (D) datasets to identify medium- and high-confidence interactors. Plotted are log2 TMT intensity fold changes for proteins between nsp2 bait channels and GFP mock transfections versus -log10 adjusted p-values. Curves for the variable cutoffs used to define high-confidence (red) or medium confidence (blue) interactors are shown. 1σ = 0.5 for (C), 1σ = 0.43 for (D). (E) Venn diagram comparing high-confidence interactors between nsp2 homologs. Sixteen unique proteins were identified each, while four proteins overlapped both data sets (listed in adjacent table).

We first determined interactors of the individual nsp2 homologs by comparing the log-transformed TMT intensity differences for prey proteins between bait and GFP samples (Figure 2C-D). We optimized variable cutoffs for high- and medium-confidence interactors based on their magnitude of enrichment compared to the GFP samples and confidence as defined by adjusted p-values (Figure 2C-D, Figure S2B-C). Using the most stringent cutoff, we identified 6 and 11 high-confidence interactors for SARS-CoV-2 and SARS-CoV-1 nsp2 respectively (Figure 2C-D). Including medium-confidence interactors, we identified 20 nsp2 interactors for each homolog, including four overlapping proteins, ERLIN1, ERLIN2, RNF170, and TMEM199 (Figure 2E).

Gene enrichment analysis shows nsp2 interactors are involved in a number of host cell processes, including metabolic processing and transport (Figure S3A). A number of these interactors are membrane-associated proteins in the ER and nucleus (Figure S3B). Detailed comparisons of gene set enrichments for individual nsp2 homologs revealed several pathways preferentially enriched for SARS-CoV-1, such as mitochondrial calcium ion transport, protein deacetylation, and negative regulation of gene expression (Figure S3C). We confirmed by immunofluorescence that SARS-CoV-1 and SARS-CoV-2 nsp2 are largely localized perinuclear and co-localize partially with the ER marker PDIA4 (Figure S3D).

To validate our findings, we cross referenced our data set with previous coronavirus interactomics studies. A prior study of SARS-CoV-1 nsp2 identified 11 host interactors, five of which overlap with our SARS-CoV-1 list, including GIGYF2, PHB, PHB2, STOML2, and EIF4E2^17^. We also cross-referenced our interactors with a recently published SARS-CoV-2 interactomics data set^18^. Interestingly, we identified 18 new interactors, though several of these share secondary interactions with the proteins identified by Gordon et al. (Figure S4).

### Quantitative comparison of SARS-CoV-1 and SARS-CoV-2 interactors

Apart from determining nsp2 host cell interactors, we sought to understand to what degree interactions vary between SARS-CoV-1 and SARS-CoV-2. Our multiplexed analysis enabled direct comparison of TMT intensities between the SARS-CoV-1 and SARS-CoV-2 nsp2 Co-IPs (Figure 3A). We validated that nsp2 bait levels are largely invariable across the replicates, enabling the direct comparison of prey protein intensities (Figure S2D). We find a subset of interactors are clearly enriched for SARS-CoV-1, including GIGYF2, HDAC8, EIF432, and PHB2 (Figure 3A). In contrast, several other interactors are enriched more strongly for SARS-CoV-2, for instance FOXK1 and NR2F2.

**Figure 3.**
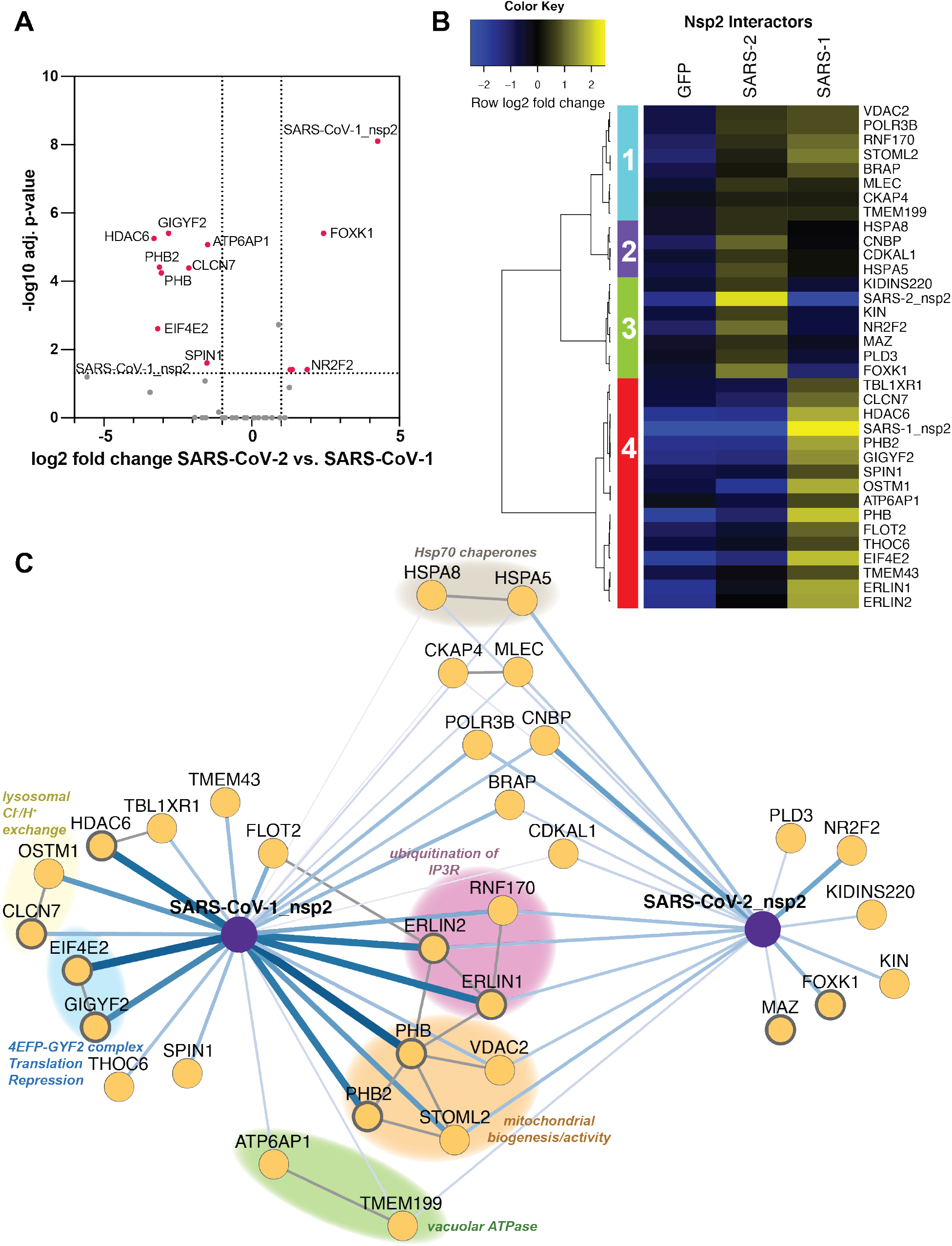
Quantitative comparison of SARS-CoV-1 and SARS-CoV-2 nsp2 interactors. (A) Volcano plot comparing interactions between nsp2 homolog from SARS-CoV-1 and SARS-CoV-2. Only high- and medium confidence interactors of nsp2 are shown. Highlighted proteins meet the filter criteria of adjusted p-value < 0.05 and | log2 fold change | >1. (B) Heatmap comparing the enrichment of SARS-CoV-1 and SARS-CoV-2 nsp2 interactors compared to GFP control. log2 fold change is color-coded and centered by row (blue low, yellow high enrichment). Hierarchical clustering using Ward’s method shown on the left was carried out on euclidean distances of log2 fold changes scaled by row. Clusters 1 and 2 corresponds to shared interactors of SARS-CoV-1 and −2 nsp2, while cluster 3 and 4 for are unique interactors for SARS-CoV-2 and SARS-CoV-1 nsp2, respectively. (C) Protein-protein interaction (PPI) network map of nsp2 homologs. Blue lines indicate viral-host PPIs, where line width corresponds to fold enrichment compared to the GFP control. Grey lines indicate annotated host-host PPIs in STRING (score > 0.75). Groups of interactors with a common functional role are highlighted.

We performed unbiased hierarchical clustering of the enrichment intensities to group the nsp2 interactors in an unbiased way. This analysis yielded four distinct clusters. Clusters 1 and 2 contained shared interactors between SARS-CoV-1 and SARS-CoV-2 nsp2. On the other hand, clusters 3 and cluster 4 contained proteins that bound exclusively to either SARS-CoV-2 or SARS-CoV-1, respectively (Figure 3B). To better visualize the relationship between shared and unique nsp2 interactors, we constructed a network plot (Figure 3C). We also included experimentally validated secondary interactions from the STRING database to group shared and unique interactors into functionally relevant subclusters.

Several of these subclusters are shared between SARS-CoV-1 and SARS-CoV-2 nsp2, for instance one including STOML2, PHB, PHB2, VDAC2. These proteins were previously shown to interact and upregulate formation of metabolically active mitochondrial membranes^26^. Another subcluster involves ERLIN1, ERLIN2, and RNF170, which form a known complex regulating ubiquitination and degradation of inositol 1,4,5-triphosphate receptors (IP_3_Rs), which in turn are channels regulating Ca^2+^ signaling from the ER to the mitochondria. Consistent with this, we detect mitochondrial calcium ion transmembrane transport as one of the unique biological processes associated with SARS-CoV-1 nsp2, but not SARS-CoV-2 (Figure S3C). Interestingly, ERLIN1 and ERLIN2 show stronger interactions with SARS-CoV-1 nsp2 than with SARS-CoV-2, whereas RNF170 interactions are similar, indicating some strain-specific preference. Additional shared interactors include a subunit of the vacuolar ATPase (ATP6AP1) and a regulatory protein (TMEM199), supporting a common role for nsp2 to influence lysosomal processes. Finally, we observe one cytosolic and one ER-resident Hsp70 chaperone (HSPA8, HSP5) as shared interactors, highlighting their role in nsp2 folding and biogenesis.

Unique SARS-CoV-2 interactors include FOXK1 and NR2F2, both of which are anti-viral transcription factors induced in response to other viruses^27,28^. We also observe an exonuclease regulator of endosomal nucleic acid sensing (PLD3)^29^, a transcription factor associated with the influenza humoral response (MAZ)^30,31^, and a DNA-binding protein implicated in B cell class switching (KIN or KIN17)^32^. In contrast, the list of unique SARS-CoV-1 interactors includes components of the 4EFP-GYF2 translation repression complex (GYGYF2, EIF4E2), lysosomal ion channels involved in chloride/proton ion exchange (CLCN7, OSTM1), and the cytosolic histone deacetylase 6 (HDAC6). While SARS-CoV-1 interactors GIGYF2 and EIF4E2 were also identified in the recent SARS-CoV-2 nsp2 dataset^18^, it is clear from our quantitative comparison that enrichment of this complex with SARS-CoV-2 nsp2 is much weaker than with SARS-CoV-1 nsp2.

### Comparative profiling of CoV nsp4 interactions

We extended our comparative analysis of host cell interactors to another CoV non-structural protein nsp4 involved in the replication complex. We applied the same AP-MS workflow used to identify nsp2 interactors (Figure 2A). In addition to SARS-CoV-1 and SARS-CoV-2 nsp4, we also included the hCoV-OC43 nsp4 construct. With this addition, we sought to probe the protein-protein interactions that differentiate strains causing severe pathogenesis versus non-severe. To this end, four co-immunoprecipitation replicates of respective nsp4 homologs were pooled into a single MS run, along with mock GFP transfected cells to differentiate non-specific background proteins (Figure 2B). The full data set included three individual MS runs, containing 40 Co-IPs (SARS-CoV-2 n = 12; SARS-CoV-1 n = 8; OC43 = 8; GFP (mock) n = 12) (Figure S5A).

As previously described, we optimized variable cutoffs for high- and medium-confidence interactors based on their magnitude enrichment compared to GFP samples (Figures 4A, S5B-C). We identified 29, 20, and 13 high-confidence interactors for SARS-CoV-2, SARS-CoV-1, and OC43 respectively using the most stringent cutoff (Figures 4A, S5B-C). Including medium-confidence interactors, we identified 86, 126, and 93 nsp4 interactors for SARS-CoV-2, SARS-CoV-1, and OC43 nsp4 homologs respectively. Comparisons of high-confidence interactors yielded 17 shared interactors between all strains (Figure 4B) or 30 medium-confidence shared interactions (Figure S5D).

**Figure 4.**
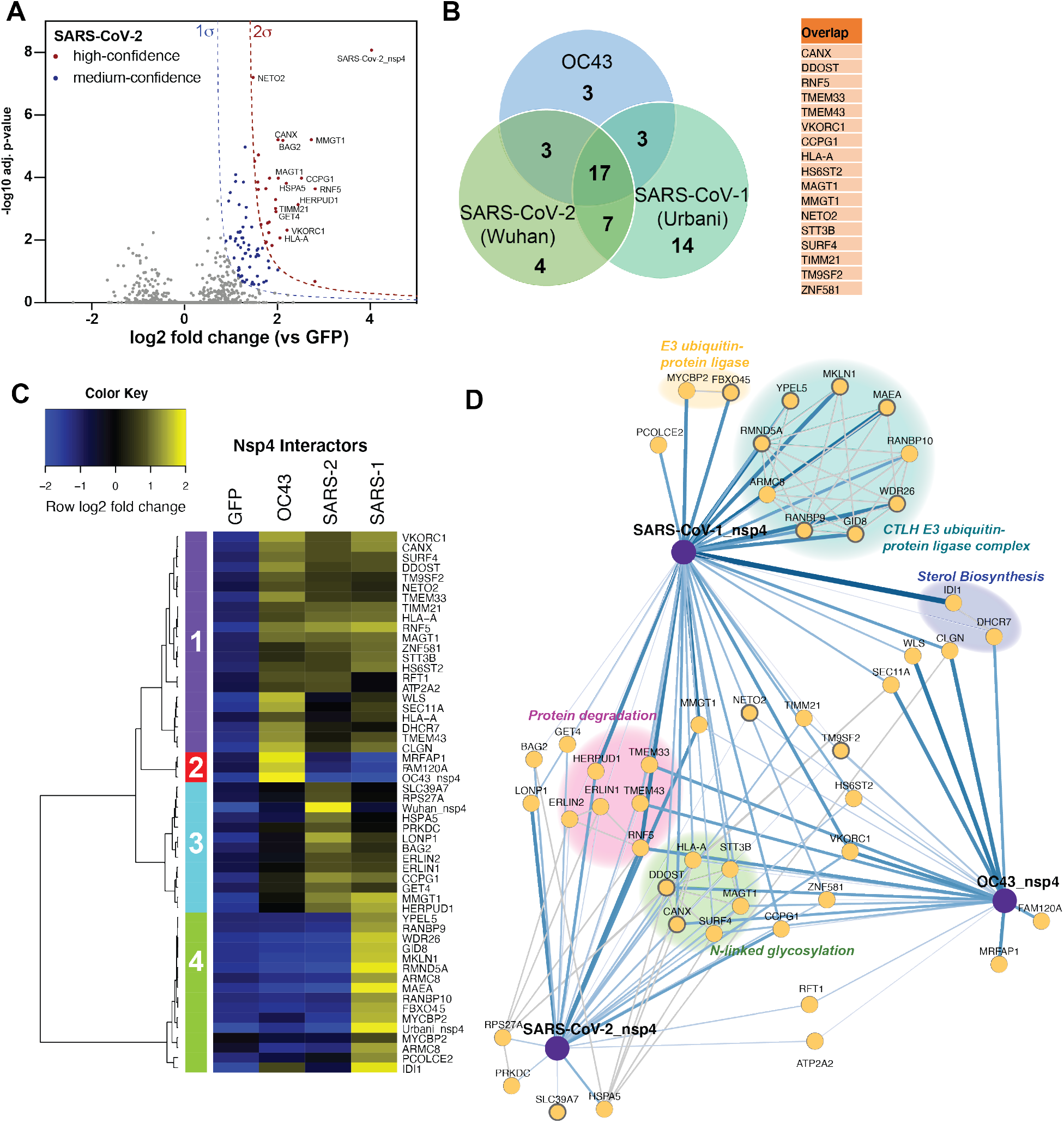
Comparative profiling of nsp4 interactions. (A) Volcano plot of the SARS-CoV-2 nsp4 datasets to identify medium- and high-confidence interactors. Plotted are log2 TMT intensity differences for proteins between nsp4 bait channels and GFP mock transfections versus -log10 adjusted p-values. Curves for the variable cutoffs used to define high-confidence (red) or medium confidence (blue) interactors are shown. 1σ = 0.66. Equivalent volcano plot for SARS-CoV-1 and OC43 nsp4 are shown in Figure S5B-C. (B) Venn diagram of interactors from nsp4 homologs. Overlapping nsp4 interactors between all strains are listed in the adjacent table. (C) Heatmap comparing the enrichment of interactors for the different nsp4 homologs. log2 fold change is color-coded and centered by row (blue low, yellow high enrichment). Hierarchical clustering using Ward’s method shown on the left was carried out on euclidean distances of log2 fold changes scaled by row. Clusters 1 corresponds to shared interactors of SARS-CoV-1, −2, and OC43 nsp4. Cluster 2, and 4 contain unique interactors for OC43 and SARS-CoV-1 nsp4, respectively, while cluster 3 contains shared interactors of SARS-CoV-1 and SARS-CoV-2. (D) Protein-protein interaction (PPI) network map of interactors of nsp4 homolog. Blue lines indicate measured viral-host PPIs, where line width corresponds to fold enrichment compared to the GFP control. Grey lines indicate annotated host-host PPIs in STRING (score > 0.75). Groups of interactors with a common functional role are highlighted.

There is relatively little overlap between our identified SARS-CoV-2 nsp4 interactors and published nsp4 interactomics data, including the recently published study of the SARS-CoV-2 interactome^18^ (Figure S6). The discrepancy could be attributed to the nsp4 constructs in our study including the N-terminal residues of nsp3, which were added to ensure proper localization and prevent the hydrophobic N-terminal region of nsp4 to serve as signal sequence^24^.

Analysis of GO-terms associated with the nsp4 interactors showed multiple enriched biological processes, such as cell organization and biogenesis, transport, and metabolic processes (Figure S7A). Interestingly, several shared SARS nsp4 interactors are associated with cell death, cellular communication, and cell differentiation. Shared interactors of all three strains are predominantly ER-membrane associated proteins, while many SARS-CoV-1 and OC43 specific interactors are annotated as nuclear-localized (Figure S7B). Comparisons of gene set enrichment analysis between strains indicate the ERAD pathway is significantly enriched for SARS strains, most strongly for SARS-CoV-1 (Figure S7C). Ubiquitin-dependent protein catabolic processes and ER mannose trimming are also strongly enriched for SARS-CoV-1. In general, processes strongly enriched for SARS-CoV-1 are less enriched for SARS-CoV-2, and to an even lesser extend for OC43.

Our multiplexed analysis of nsp4 homolog Co-IPs enabled direct comparison across strains (Figure S8A-C). We validated that nsp4 bait levels were mostly similar across replicates, allowing for direct comparison of bait protein intensities (Figure S8D). Unbiased hierarchical clustering of enrichment intensities to group nsp4 interactors yielded four distinct clusters. Clusters 1 contained common interactors of all nsp4 homologs (Figure 4C), while cluster 3 contains shared interactors of SARS-CoV-2, SARS-CoV-1 nsp4 that displayed weaker enrichment with OC43. In contrast, cluster 2 and 4 contained unique interactors enriched for OC43 and SARS-CoV-1 nsp4, respectively. To visualize functionally relevant subclusters of shared and unique nsp4 interactors, we constructed a network plot, including high-confidence interactions (score > 0.75) from the String database (Figure 4D). Inclusion of all median-confidence interactors of the nsp4 homologs yielded a similar clustering and network organization (Figures S8E, S9).

We identified several common interactors across all three nsp4 homologs. These include components of UPR signaling (TMEM33) and ER-phagy (CCPG1). We also identify RNF5, an ER-localized E3 ubiquitin ligase known to modulate anti-viral innate immune signaling^33,34^, and VKORC1, which reduces Vitamin K, a key cofactor for several coagulation factor proteins^35^. Not surprisingly, given that nsp4 is a glycosylated protein, we also identify several members of the *N*-linked glycosylation machinery (STT3B, MAGT1, CANX, DDOST) (Figure 4D) in all three strains.

We identified several shared interactors between SARS-CoV-1 and SARS-CoV-2 that were absent in OC43. These include the ERLIN1/2 complex, LONP1, HERPUD1, GET4, and BAG2, all of which are involved in a facet of ER homeostasis, proteostasis or trafficking (Figure 4D). The ERLIN1/2 complex was also identified in the nsp2 data set (Figure 3B-C) and shows comparable enrichment values between SARS-CoV-1 and SARS-CoV-2. Interestingly, the other four overlapping interactors all exhibit increased enrichment for SARS-CoV-2 versus SARS-CoV-1. LONP1 is a mitochondrial peptidase responsible for removing the majority of damaged mitochondrial proteins via proteolysis. The unfolded protein response (UPR) induces HERPUD1 expression, which is involved in the ER-associated degradation pathway (ERAD) to maintain ER homeostasis ^36^. BAG2 serves as a co-chaperone for HSP70 chaperones, acting as a nucleotide exchange factor to regulate chaperone-client interactions through modulating HSP70 ATPase rates ^37^, while GET4 is part of a complex driving trafficking of tail-anchored proteins to the ER ^38^.

We observed shared interactors between OC43 and SARS-CoV-2, such as the *N*-glycosylation factor RFT1 ^39,40^ and a sarcoplasmic/endoplasmic reticulum calcium ATPase (SERCA – ATP2A2) ^41^. In addition, we identified shared interactors between OC43 and SARS-CoV-1, including a regulator of UPR-mediated apoptosis (WLS or GPR177) ^42^, a member of the signal peptidase complex (SEC11A) ^43^, and factors involved in cholesterol synthesis (IDI1, DHCR7) ^44–47^. WLS, SEC11A, and DHRC7 exhibited higher enrichment for OC43, whereas IDI1 was more greatly enriched for SARS-CoV-1. Consistent with this observation, we identified sterol metabolic process as one of the unique processes enriched for OC43 nsp4.

In addition to shared interactors, we found several unique interactors for SARS-CoV-2, including the monoubiquitin-ribosomal fusion protein (RPS27A), a Golgi/ER-resident zinc receptor that has been shown to regulate TNF receptor trafficking and necroptosis (SLC39A7), and the ER-resident Hsp70 chaperone BiP (HSPA5). The latter two play distinct roles in regulating ER homeostasis and proteostasis. In contrast, only two unique OC43 nsp4 interactors were identified: a target of the NEDD8-Cullin E3 ligase pathway (MRFAP1)^48^ and FAM120A, an RNA-binding protein found to serve as a scaffolding protein for the IL13 signaling pathway (Figure S8B-C) ^49,50^. Both of these proteins are localized to the nucleus (Figure S7B). Lastly, we identified a large cluster of unique SARS-CoV-1 nsp4 interactors that compose the CTLH E3 ubiquitin ligase complex (Figure 4D). This nuclear complex maintains cell proliferation rates, likely through the ubiquitination of the transcription factor Hbp1, a negative regulator of cell proliferation^51^. This complex is highly enriched for SARS-CoV-1 specifically, presenting one of the most profound differences in interaction profile (Figure S8A,C).

The fact that both OC43 and SARS-CoV-1 nsp4 displayed prominent interactions with nuclear proteins prompted us to evaluate the cellular localization of the protein by immunofluorescence. We detected perinuclear puncti for all constructs which partially co-localized with the ER marker PDIA4 (Figure S10), consistent with prior studies^22^. However, for SARS-CoV-1 and OC43 nsp4, we also detected measurable signal in the nucleus, supporting a nuclear function and the observed interactions with protein in the nucleus.

### Enrichment of mitochondria-associated membrane proteins as nsp2 and nsp4 interactors

In our evaluation of cellular compartment GO-terms, we noticed that nsp2 and nsp4 interactors are enriched in membranes of the endoplasmic reticulum and the mitochondria (Figures S3B, S7B). In particular, ERLIN1/2 and RNF170 form an E3 ubiquitin ligase complex known to localize to the interface between the ER and mitochondria, regions termed mitochondria-associated membranes (MAMs). We therefore probed our data set for any other MAMs-associated nsp2 and nsp4 interactors. We cross-referenced our interactor lists with three published data sets that specifically characterized the MAMs proteome^52–54^ and identified 17 proteins associated with MAMs (Figure 5A-B). Seven of these factors solely interact with nsp2, eight proteins solely interact with one or more strains of nsp4, and the ERLIN1/2 complex interacts with both nsp2 and nsp4 (Figure 3C, 4D). Interestingly, the ERLIN1/2 complex only interacts with SARS-CoV-1 and −2 proteins and not OC43. SARS-CoVs may use ERLIN1/2 to regulate ER Ca^2+^ signaling and the myriad of downstream host processes controlled by this signaling pathway (Figure 5C).

**Figure 5.**
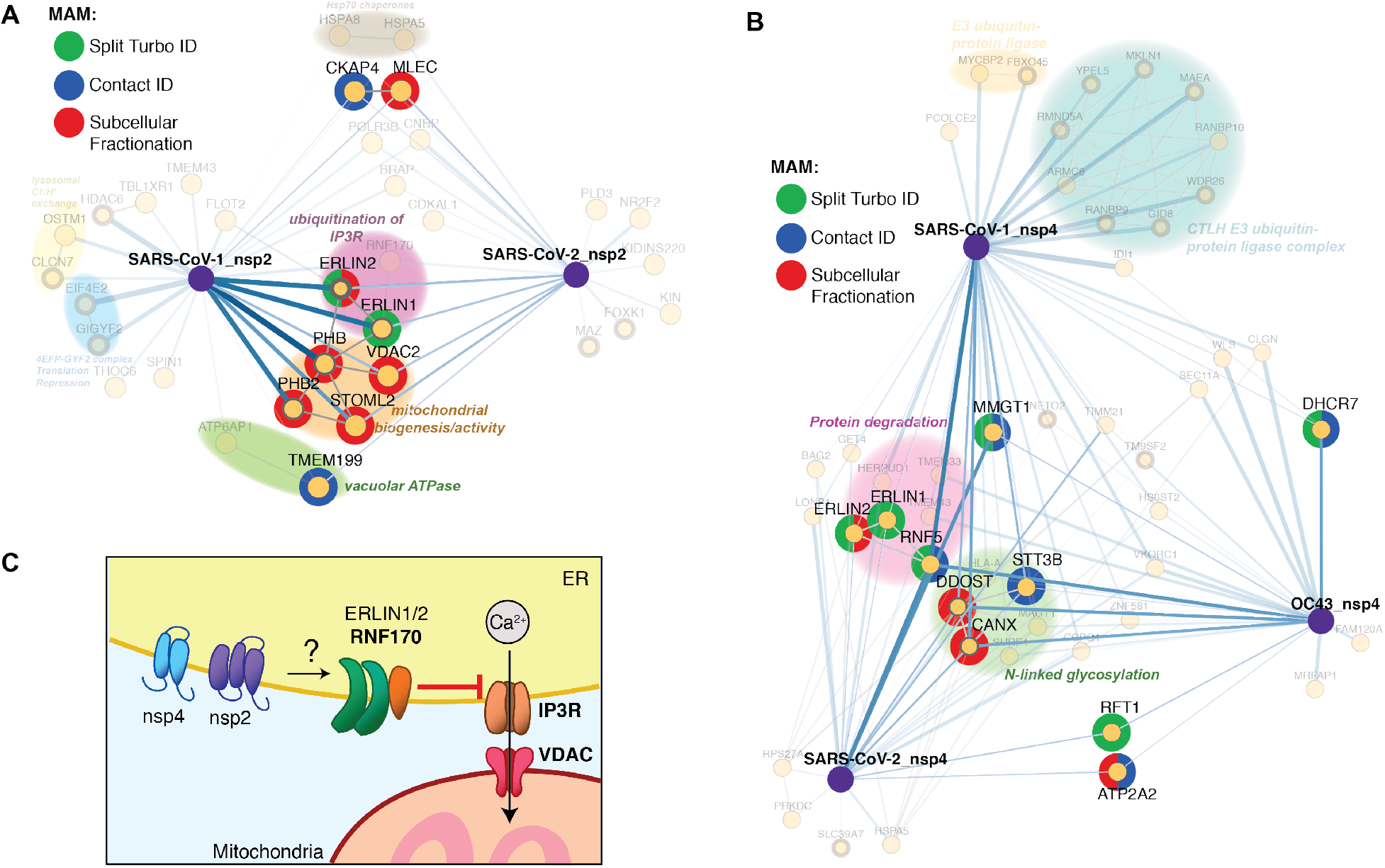
Enrichment of mitochondria-associated membrane (MAM) proteins as nsp2 and nsp4 interactors. (A–B) Interactors of nsp2 (A) and nsp4 (B) homolog annotated for MAM proteins. The lists of interactors were cross-referenced with previous publications profiling the MAM proteome (Split-Turbo ID^52^, Contact-ID^53^, and subcellular fractionation^54^). (C) Proposed model for how SARS-CoV nsp2 and nsp4 utilize ERLIN1/2 and interacting protein factors to regulate ER Ca^2+^ signaling at MAMs.

## DISCUSSION

Our analysis enables both the identification of interactors for SARS-CoV-1, SARS-CoV-2, and OC43 homologs of nsp2 and nsp4, and comparative quantitative enrichment to differentiate between shared and unique host cell binding partners. We acknowledge the limitations of using transiently transfected viral proteins for AP-MS. Viral infection results in a collection of both protein-protein and RNA-protein interactions and our approach cannot account for how these events influence the interactions of nsp2 and nsp4. However, given the logistical barriers to handling BSL-3 viruses, paired with the urgency of the current pandemic, our workflow is an efficient system to perform comparative analysis and generate a shortlist of interactors to prioritize for further investigation.

We identify several nsp2 interactors shared across SARS strains, including STOML2, and prohibitins (PHB and PHB2), which were previously identified as interacting with SARS-CoV-1^17^. These proteins work in tandem to induce formation of metabolically active mitochondrial membranes to regulate mitochondrial biogenesis. Increased levels of STOML2 are associated with increased ATP production and reduced apoptosis induction^26^. This conserved interaction for SARS strains presents an avenue for nsp2 to increase mitochondrial metabolism and stall apoptosis to maintain a pro-viral cellular environment. Additionally, STOML2 has been found to play a key role in stabilizing hepatitis C virus replication complexes^55^ and PHB has been shown to promote entry of both Chikungunya virus^56^ and enterovirus 71^57^. These factors may prove effective pan-RNA virus targets for host-directed therapies.

For nsp4, we identify multiple unique SARS-CoV-1 interactors, most notably members of the CTLH E3 ubiquitin ligase complex. This complex is known to regulate levels of Hbp1, a negative regulator of proliferative genes^51^ and was previously shown to interact with the dengue viral protein NS2B3^15^, implicating this complex as a target for RNA viruses to influence cell proliferation. We also identified the FBXO45-MYCBP2 E3 ubiquitin ligase complex, which has been shown to prevent cell death in mitosis ^58^. Together, this may support a role in SARS-CoV-1 nsp4 co-opting host ubiquitin complexes to extend cell viability during infection to promote viral replication.

Furthermore, we find components of the cholesterol biosynthesis pathway, IDI1 and DHCR7, which were specifically enriched for SARS-CoV-1 and OC43 nsp4 respectively. IDI1 has been shown to be downregulated by host cells in response to CMV-infection-induced interferons^44^ and is upregulated by both HIV and HCV during infection^45,46^. DHCR7 is downregulated during RNA virus infection in macrophages to promote IRF3 signaling and IFN-1 production. Moreover, inhibition of DHCR7 aids in clearance of multiple RNA viruses^47^. These previous findings indicate that interactions with IDI1 and DHCR7 may provide means for coronaviruses to counteract anti-viral responses. Interestingly, these interactions with the aforementioned E3 ligase complexes and cholesterol biogenesis factors are not enriched for SARS-CoV-2 nsp4, implying that SARS-CoV-2 pathogenesis may not require these interactions.

As a whole, it appears that SARS-CoV-2 homologs differ from SARS-CoV-1 not by gaining new interactions, but rather by losing network nodes. This is emphasized in the gene enrichment analysis of nsp2 and nsp4 (Figure S3C, S7C), in which multiple pathways are more strongly enriched for SARS-CoV-1, as well as in the nsp4 interactome (Figure 4C-D), particularly with the absence of E3 ligase complex interactions for SARS-CoV-2. It will be important to investigate potential functional implications of the engagement of the E3 ubiquitin ligase, as well innate immune signaling factors on CoV infections and the course of pathogenicity for the divergent strains.

A particularly noteworthy finding is the identification of 17 MAMs factors in the combined nsp2 and nsp4 datasets, based on cross-referencing interactors with previously published proteomics studies of MAMs proteins^52–54^. Given the prominence of these interactions, it is tempting to speculate that nsp2 and nsp4 localize to MAMs and influence processes at these important organelle contact sites (Figure 5C). MAMs are nodes for innate immune signaling and apoptosis pathways, both of which are common targets for viral manipulation.

In particular, we identify the E3 ubiquitin ligase RNF5 interacting with all nsp4 homologs. RNF5 targets STING for degradation, which stabilizes retinoic acid-inducible gene-I (RIG-I) and mitochondrial antiviral-signaling protein (MAVS) interactions at MAMs, thereby inducing inteferon-1 and −3 production via IRF3 and NF-κB signaling^33,34^. RIG-I is one of the main viral RNA genome sensors in host cells; therefore, it is possible that nsp4 increases targeting of RNF5 to MAMs to inhibit downstream signaling of RIG-1.

We also identify the ERLIN1/2 complex in both nsp2 and nsp4 data sets. In the nsp2 interaction network, the complex is associated with a different E3 ligase, RNF170. RNF170 has been shown to inhibit innate immune signaling by targeting TLR3 for degradation, thereby blocking IRF3 and NF-κB signaling pathways^59^. In addition, ERLIN1/2 acts in concert with RNF170 to target the inositol-1,4,5-triphophate receptor (IP_3_R) for degradation via polyubiquination^60^. IP_3_R is an ER-resident Ca^2+^ channel integral in the formation of MAMs^61,62^. Calcium flux at MAMs has been shown to increase mitochondrial calcium uptake, which increases ATP production, thereby benefitting active viral replication^63^. Indeed, several other viruses have been shown to influence ER Ca^2+^ exchange, such as the rotavirus protein NSP4 and the HIV-1 protein Nef, which both cause depletion of ER Ca^2+^ stores through modulation of IP_3_R activity^64^. Another prominent example includes the human cytomegalovirus protein vMIA, which increases ER Ca^2+^ export at MAMs through IP_3_R into the mitochondria ^63^. Previous studies have shown the SARS-CoV-1 E protein acts as a channel to leak ER calcium stores during infection^65^, but to our knowledge, no such features have been attributed to either nsp2 or nsp4. Thus, manipulation of ER Ca^2+^ signaling via IP_3_R regulation may represent a novel method by which coronaviruses manipulate mitochondrial function (Figure 5C). In support of this, a recent study found that IP_3_R is significantly reduced during SARS-CoV-2 infection^66^. Further studies will be important to evaluate whether ER calcium exchange and mitochondrial metabolism could impact coronavirus infection.

## METHODS

### Protein Expression constructs

Coding sequences for nsp2 and nsp4 were obtained from GenBank (MN908947 SARS-CoV-2 isolate Wuhan-Hu-1; AY278741 SARS-CoV-1 Urbani; NC_006213 hCoV OC43 strain ATCC VR-759). Human codon optimized sequences were designed, genes synthesized, and cloned into pcDNA3.1-(+)-C-DYK (nsp4) to append a C-terminal FLAG tag, or into pcDNA3.1-(+)-N-DYK (nsp2) to append an N-terminal FLAG tag (GenScript).

### Cell Culture and Transfection

HEK293T cells were maintained in Dulbecco’s Modified Eagle’s Medium (DMEM) with high glucose and supplemented with 10% fetal bovine serum (FBS), 1% penicillin/streptomycin, and 1% glutamine. Cells were kept at 37°C, 5% CO2. Generally, 2 × 10^6^ cells were seeded into 10cm dishes. 24 hours post-seeding, cells were transfected with 5μg nsp2, nsp4, or fluorescent control DNA constructs in pcDNA3.1-(+)-C/N-DYK vectors using a calcium phosphate method. Media was exchanged 16 hours post-transfection, and cells were harvested 24 hours after changing media.

### Immunoprecipitation

Cells were collected and washed with PBS. Immunoprecipitation samples were lysed by resuspension in TNI buffer (50 mM Tris pH 7.5, 150mM NaCl, 0.5% IGEPAL-CA-630) with Roche cOmplete protease inhibitor on ice for at least 10 minutes, followed by sonication in a room temperature water bath for 10 minutes. Lysates were cleared by centrifugation at 17,000 x g for 10-20 minutes. Sepharose 4B resin (Sigma) and G1 anti-DYKDDDDK resin (GenScript) were pre-washed 4x with the respective lysis buffer for each sample. Protein concentrations in cleared lysates were normalized using BioRad Protein Assay Dye and added to 15 μL Sepharose 4B resin for 1 hour, rocking at 4°C. Resin was collected by centrifugation for 5-10 minutes at 400 x g and pre-cleared supernatant was added directly to 15 μL G1 anti-DYKDDDDK resin and rocked at 4°C overnight. The next day, supernatant was removed and resin was washed 4 times with the respective lysis buffer. Bound proteins were eluted with the addition of modified 3x Laemelli buffer (62.5 mM Tris, 6% SDS) for 30 minutes at room temperature followed by 15 minutes at 37°C, followed by a second elution for 5-15 minutes at 37°C. 10% of elution was set aside for SDS-PAGE and silver staining to confirm immunoprecipitation efficiency, and the remainder was prepared for mass spectrometry. Silver staining was performed using a Pierce Silver Stain kit (Thermo Scientific).

### Tandem Mass Tag Sample Preparation

Sample preparation was carried out as described^67^. Briefly, eluted proteins were precipitated in methanol/chloroform/water (3:1:3), washed twice in methanol, and protein pellets were air dried. Pellets were resuspended in 1% Rapigest SF (Waters), reduced, and alkylated. Proteins were digested in trypsin-LysC overnight. Digested peptides were labeled using 11-plex or 16-plex tandem mass tag (TMT or TMTpro) reagents (Thermo Scientific), pooled, and acidified using formic acid. Cleaved Rapigest was removed by centrifugation of samples at 17,000 x g for 30 min.

### MudPIT liquid chromatography-mass spectrometry Analysis

Triphasic MudPIT microcolumn were prepared as described^68^. Individual pooled TMT proteomics samples were directly loaded onto the microcolumns using a high-pressure chamber followed by a wash with 5% acetonitrile, 0.1% formic acid in water (v/v) for 30 min. Peptides were analyzed by liquid chromatography-mass spectrometry on an Exploris 480 in line with an Ultimate 3000 nanoLC system (Thermo Fisher). The MudPIT microcolumns were installed on a column switching valve on the nanoLC systems followed by 20 cm fused silica microcapillary column (ID 100μm) ending in a laser-pulled tip filled with Aqua C18, 3μm, 100 Å resin (Phenomenex). MudPIT runs were carried out by 10μL sequential injection of 0, 10, 20, 40, 60, 80, 100 % buffer C (500mM ammonium acetate, 94.9% water, 5% acetonitrile, 0.1% formic acid), followed by a final injection of 90% C, 10% buffer B (99.9% acetonitrile, 0.1% formic acid v/v). Each injection was followed by a 130 min gradient using a flow rate of 500nL/min (0 – 6 min: 2% buffer B, 8 min: 5% B, 100 min: 35% B, 105min: 65% B, 106 – 113 min: 85% B, 113 – 130 min: 2% B). Electrospray ionization was performed directly from the tip of the microcapillary column using a spray voltage of 2.2 kV, ion transfer tube temperature of 275°C and RF Lens of 40%. MS1 spectra were collected using the following settings: scan range 400 – 1600 m/z, 120,000 resolution, AGC target 300%, and automatic injection times. Data-dependent tandem mass spectra were obtained using the following settings: monoisotopic peak selection mode: peptide, included charge state 2 – 7, TopSpeed method (3s cycle time), isolation window 0.4 m/z, HCD fragmentation using a normalized collision energy of 32, resolution 45,000, AGC target 200%, automatic injection times, and dynamic exclusion (20 ppm window) set to 60 s.

### Experimental layout and data analysis

The nsp2 AP-MS experiments included three individual MS runs combining 34 Co-AP samples (SARS-CoV-2 n = 13; SARS-CoV-1 n = 9; GFP (mock) n = 12). Samples were distributed to TMTpro 16plex or TMT11plex channels as outlined in Figure S2A. The nsp4 AP-MS experiments consisted of three individual MS runs, containing 40 Co-IPs (SARS-CoV-2 n = 12; SARS-CoV-1 n = 8; OC43 = 8; GFP (mock) n = 12). Samples were distributed to TMTpro 16plex channels as outlined in Figure S5A. Identification and quantification of peptides and proteins were carried out in Proteome Discoverer 2.4 (Thermo Fisher) using a SwissProt human database (Tax ID 9606, release date 11/23/2019). CoV nsp2 and nsp4 protein sequences were added manually. Searches were conducted in Sequest HT using the following setting: Trypsin cleavage with max. 2 missed cleavage sites, minimum peptide length 6, precursor mass tolerance 20 ppm, fragment mass tolerance 0.02 Da, dynamic modifications: Met oxidation (+15.995 Da), Protein N-terminal Met loss (−131.040 Da), Protein N-terminal acetylation (+42.011 Da), static modifications: Cys carbamidomethylation (+57.021 Da), TMTpro or TMT6plex at Lys and N-termini (+304.207 Da for TMTpro or +229.163 for TMT6plex). Peptide IDs were filtered using the Percolator node using an FDR target of 0.01. Proteins were filtered based on a 0.01 FDR requiring two peptide IDs per protein, and protein groups were created according to a strict parsimony principle. TMT reporter ions were quantified using the reporter ion quantification considering unique and razor peptides, and excluding peptides with co-isolation interference greater than 25%. Peptide abundances were normalized based on total peptide amounts in each channel assuming similar levels of background signal in the APs. Protein quantification roll-up used all quantified peptides. Pairwise ratios between conditions were calculated based on total protein abundances and ANOVA on individual proteins was used to test for changes in abundances and to report adjusted p-values. To filter high-confidence interactors of individual CoV nsp proteins, we used a variable filter combining log2 fold enrichment and adjusted p-value according to a published method^69^. Briefly, the histogram of log2 protein abundance fold changes between nsp-transfected versus mock-transfected groups were fitted to a gaussian curve using a nonlinear least square fit to determine the standard deviation σ (see Figure S2B-C). Fold change cutoffs for high-confidence and medium-confidence interactors were based on 2 σ, or 1 σ, respectively. For actual cutoffs taking into consideration adjusted p-values, we utilized a hyperbolic curve *y > c / (x – x_0_)*, where *y* is the adj. p-value, *x* the log2 fold change, *x*_*0*_ corresponds to the standard deviation cutoff (2 σ or 1 σ), and *c* is the curvature (c = 0.4 for 1 σ, and 0.8 for 2 σ). (see Figures 2C-D, 4A, S5B-C).

### Gene set enrichment analysis

GO-term categories for biological processes and cellular components for interactors were based on assignment in the Proteome Discoverer Protein Annotation node. Gene set enrichment analysis was conducted in EnrichR^70^. The analysis was conducted separately for sets of interactors of individual nsp2 or nsp4 homologs, and GO-terms for biological processes were filtered by adjusted p-values < 0.1. Redundant GO-terms were grouped manually based on overlapping genes in related terms.

### Network plots and identification of overlapping interactions with published data

Extended and overlapping interactomes between novel interactors identified in this study and previously published interactors^18^ were generated by scraping the top n interactors of each primary prey protein on the STRING database using the python API. We established an extended secondary interactome by searching for the top 20 and top 30 STRING db interactors of the nsp4 primary interactors and nsp2 interactors respectively using limit parameter in STRING API and searching against the human proteome (species 9606). We then compared the extended interactomes of our data with the previously published data by dropping any secondary interactors that did not appear in both data sets. Next, we concatenated the primary interactors from our data, the primary interactors from the published data, and the overlapping secondary interactors into a single data frame. Finally, we searched the overlapping secondary interactors against the STRING database human proteome to determine interactors between secondary interactors with a threshold of greater than 50% likelihood in the experimental score category. The results were plotted in Cytoscape.

### Immunofluorescence Confocal Microscopy

HEK293T cells were cultured on glass-bottom culture dishes (MatTek, P35G-0-14-C) and transfected with CoV expression constructs as previously described. Cells were fixed with 4% paraformaldehyde-PBS, washed thrice with PBS, then permeabilized in 0.2% Triton-X (in PBS). After three PBS washes, cells were blocked in PBS with 1% BSA with 0.1% Saponin (blocking buffer). After blocking, cells were incubated with anti-PDIA4 primary antibody (Protein Tech, 14712-1-AP) in blocking buffer (1:1000 dilution) for 1 hour at 37°C. After three PBS washes, cells were incubated with AlexFluor 488-conjugated anti-rabbit goat antibody (ThermoFisher, A-11008) in blocking buffer (1:500 dilution) at room temperature for 30 min. Cells were then stained with M2 FLAG primary antibody (SigmaAldrich, F1804) and AlexFluor 594-conjugated anti-mouse goat antibody (ThermoFisher, A-11005) using the same conditions. Cells were then mounted in Prolong Gold with DAPI stain (ThermoFisher, P36935). Cells were imaged using an LSM-880 confocal microscope (Zeiss) and images were merged using Image J software.

## ACKNOWLEDGMENTS

We thank members of the Plate group for critical reading of the manuscript. This work was funded by T32 AI112541, and the National Science Foundation Graduate Research Fellowship Program (K.M.A); T32 GM008554 (J.P.D), T32 GM065086 (E.F.M.), NIGMS R35 award (1R35GM133552), as well as Vanderbilt University startup funds.

## REFERENCES

(1) Fehr, A. R.; Perlman, S. Coronaviruses: An Overview of Their Replication and Pathogenesis. Methods Mol. Biol. 2015, 1282, 1. https://doi.org/10.1007/978-1-4939-2438-7_1.

(2) Novel Coronavirus (2019-NCoV) Situation Report 1; 2020.

(3) Hoffmann, M.; Kleine-Weber, H.; Schroeder, S.; Mü, M. A.; Drosten, C.; Pö, S. SARS-CoV-2 Cell Entry Depends on ACE2 and TMPRSS2 and Is Blocked by a Clinically Proven Protease Inhibitor. Cell 2020, 181, 271–280. https://doi.org/10.1016/j.cell.2020.02.052.

(4) Perlman, S.; Netland, J. Coronaviruses Post-SARS: Update on Replication and Pathogenesis. Nature Reviews Microbiology. 2009, pp 439–450. https://doi.org/10.1038/nrmicro2147.

(5) Blanco-Melo, D.; Nilsson-Payant, B. E.; Liu, W.-C.; Lim, J. K.; Albrecht, R. A.; Tenoever, B. R. SARS-CoV-2 Infection Induces Low IFN-I and-III Levels with a Moderate ISG Response. Cell 2020. https://doi.org/10.1016/j.cell.2020.04.026.

(6) Fung, T. S.; Liu, D. X. Coronavirus Infection, ER Stress, Apoptosis and Innate Immunity. Frontiers in Microbiology. Frontiers Research Foundation June 17, 2014, p 296. https://doi.org/10.3389/fmicb.2014.00296.

(7) Tan, Y.-X.; Tan, T. H. P.; Lee, M. J.-R.; Tham, P.-Y.; Gunalan, V.; Druce, J.; Birch, C.; Catton, M.; Fu, N. Y.; Yu, V. C.; Tan, Y.-J. Induction of Apoptosis by the Severe Acute Respiratory Syndrome Coronavirus 7a Protein Is Dependent on Its Interaction with the Bcl-XL Protein. J. Virol. 2007, 81 (12), 6346–6355. https://doi.org/10.1128/jvi.00090-07.

(8) Yeung, M. L.; Yao, Y.; Jia, L.; Chan, J. F. W.; Chan, K. H.; Cheung, K. F.; Chen, H.; Poon, V. K. M.; Tsang, A. K. L.; To, K. K. W.; Yiu, M. K.; Teng, J. L. L.; Chu, H.; Zhou, J.; Zhang, Q.; Deng, W.; Lau, S. K. P.; Lau, J. Y. N.; Woo, P. C. Y.; Chan, T. M.; Yung, S.; Zheng, B. J.; Jin, D. Y.; Mathieson, P. W.; Qin, C.; Yuen, K. Y. MERS Coronavirus Induces Apoptosis in Kidney and Lung by Upregulating Smad7 and FGF2. Nat. Microbiol. 2016, 1 (3), 1–8. https://doi.org/10.1038/nmicrobiol.2016.4.

(9) Yue, Y.; Nabar, N. R.; Shi, C. S.; Kamenyeva, O.; Xiao, X.; Hwang, I. Y.; Wang, M.; Kehrl, J. H. SARS-Coronavirus Open Reading Frame-3a Drives Multimodal Necrotic Cell Death. Cell Death Dis. 2018, 9 (9), 1–15. https://doi.org/10.1038/s41419-018-0917-y.

(10) von Brunn, A.; Teepe, C.; Simpson, J. C.; Pepperkok, R.; Friedel, C. C.; Zimmer, R.; Roberts, R.; Baric, R.; Haas, J. Analysis of Intraviral Protein-Protein Interactions of the SARS Coronavirus ORFeome. PLoS One 2007, 2 (5), e459. https://doi.org/10.1371/journal.pone.0000459.

(11) Pfefferle, S.; Schö Pf, J.; Kö Gl, M.; Friedel, C. C.; Mü Ller, M. A.; Carbajo-Lozoya, J.; Stellberger, T.; Von Dall’armi, E.; Herzog, P.; Kallies, S.; Niemeyer, D.; Ditt, V.; Kuri, T.; Zü St, R.; Pumpor, K.; Hilgenfeld, R.; Schwarz, F.; Zimmer, R.; Steffen, I.; Weber, F.; Thiel, V.; Herrler, G.; Thiel, H.-J. R.; Schwegmann-Weßels, C.; Pö Hlmann 10, S.; Rgen Haas, J.; Drosten, C.; Von Brunn, A. The SARS-Coronavirus-Host Interactome: Identification of Cyclophilins as Target for Pan-Coronavirus Inhibitors. 2011. https://doi.org/10.1371/journal.ppat.1002331.

(12) V’kovski, P.; Gerber, M.; Kelly, J.; Pfaender, S.; Ebert, N.; Braga Lagache, S.; Simillion, C.; Portmann, J.; Stalder, H.; Gaschen, V.; Bruggmann, R.; Stoffel, M. H.; Heller, M.; Dijkman, R.; Thiel, V. Determination of Host Proteins Composing the Microenvironment of Coronavirus Replicase Complexes by Proximity-Labeling. Elife 2019, 8. https://doi.org/10.7554/eLife.42037.

(13) Jean Beltran, P. M.; Cook, K. C.; Cristea, I. M. Exploring and Exploiting Proteome Organization during Viral Infection. J. Virol. 2017, 91 (18). https://doi.org/10.1128/jvi.00268-17.

(14) Hashimoto, Y.; Sheng, X.; Murray-Nerger, L. A.; Cristea, I. M. Temporal Dynamics of Protein Complex Formation and Dissociation during Human Cytomegalovirus Infection. Nat. Commun. 2020, 11 (1). https://doi.org/10.1038/s41467-020-14586-5.

(15) Shah, P. S.; Link, N.; Jang, G. M.; Sharp, P. P.; Zhu, T.; Swaney, D. L.; Johnson, J. R.; Von Dollen, J.; Ramage, H. R.; Satkamp, L.; Newton, B.; Hüttenhain, R.; Petit, M. J.; Baum, T.; Everitt, A.; Laufman, O.; Tassetto, M.; Shales, M.; Stevenson, E.; Iglesias, G. N.; Shokat, L.; Tripathi, S.; Balasubramaniam, V.; Webb, L. G.; Aguirre, S.; Willsey, A. J.; Garcia-Sastre, A.; Pollard, K. S.; Cherry, S.; Gamarnik, A. V.; Marazzi, I.; Taunton, J.; Fernandez-Sesma, A.; Bellen, H. J.; Andino, R.; Krogan, N. J. Comparative Flavivirus-Host Protein Interaction Mapping Reveals Mechanisms of Dengue and Zika Virus Pathogenesis. Cell 2018, 175 (7), 1931–1945.e18. https://doi.org/10.1016/j.cell.2018.11.028.

(16) Nicod, C.; Banaei-Esfahani, A.; Collins, B. C. Elucidation of Host–Pathogen Protein– Protein Interactions to Uncover Mechanisms of Host Cell Rewiring. Curr. Opin. Microbiol. 2017, 39, 7–15. https://doi.org/10.1016/j.mib.2017.07.005.

(17) Cornillez-Ty, C. T.; Liao, L.; Yates, J. R.; III; Kuhn, P.; Buchmeier, M. J. Severe Acute Respiratory Syndrome Coronavirus Nonstructural Protein 2 Interacts with a Host Protein Complex Involved in Mitochondrial Biogenesis and Intracellular Signaling. J. Virol. 2009, 83 (19), 10314. https://doi.org/10.1128/JVI.00842-09.

(18) Gordon, D. E.; Jang, G. M.; Bouhaddou, M.; Xu, J.; Obernier, K.; White, K. M.; O’Meara, M. J.; Rezelj, V. V.; Guo, J. Z.; Swaney, D. L.; Tummino, T. A.; Huettenhain, R.; Kaake, R. M.; Richards, A. L.; Tutuncuoglu, B.; Foussard, H.; Batra, J.; Haas, K.; Modak, M.; Kim, M.; Haas, P.; Polacco, B. J.; Braberg, H.; Fabius, J. M.; Eckhardt, M.; Soucheray, M.; Bennett, M. J.; Cakir, M.; McGregor, M. J.; Li, Q.; Meyer, B.; Roesch, F.; Vallet, T.; Mac Kain, A.; Miorin, L.; Moreno, E.; Naing, Z. Z. C.; Zhou, Y.; Peng, S.; Shi, Y.; Zhang, Z.; Shen, W.; Kirby, I. T.; Melnyk, J. E.; Chorba, J. S.; Lou, K.; Dai, S. A.; Barrio-Hernandez, I.; Memon, D.; Hernandez-Armenta, C.; Lyu, J.; Mathy, C. J. P.; Perica, T.; Pilla, K. B.; Ganesan, S. J.; Saltzberg, D. J.; Rakesh, R.; Liu, X.; Rosenthal, S. B.; Calviello, L.; Venkataramanan, S.; Liboy-Lugo, J.; Lin, Y.; Huang, X. P.; Liu, Y. F.; Wankowicz, S. A.; Bohn, M.; Safari, M.; Ugur, F. S.; Koh, C.; Savar, N. S.; Tran, Q. D.; Shengjuler, D.; Fletcher, S. J.; O’Neal, M. C.; Cai, Y.; Chang, J. C. J.; Broadhurst, D. J.; Klippsten, S.; Sharp, P. P.; Wenzell, N. A.; Kuzuoglu, D.; Wang, H. Y.; Trenker, R.; Young, J. M.; Cavero, D. A.; Hiatt, J.; Roth, T. L.; Rathore, U.; Subramanian, A.; Noack, J.; Hubert, M.; Stroud, R. M.; Frankel, A. D.; Rosenberg, O. S.; Verba, K. A.; Agard, D. A.; Ott, M.; Emerman, M.; Jura, N.; von Zastrow, M.; Verdin, E.; Ashworth, A.; Schwartz, O.; D’Enfert, C.; Mukherjee, S.; Jacobson, M.; Malik, H. S.; Fujimori, D. G.; Ideker, T.; Craik, C. S.; Floor, S. N.; Fraser, J. S.; Gross, J. D.; Sali, A.; Roth, B. L.; Ruggero, D.; Taunton, J.; Kortemme, T.; Beltrao, P.; Vignuzzi, M.; García-Sastre, A.; Shokat, K. M.; Shoichet, B. K.; Krogan, N. J. A SARS-CoV-2 Protein Interaction Map Reveals Targets for Drug Repurposing. Nature 2020. https://doi.org/10.1038/s41586-020-2286-9.

(19) Brito, A. F.; Pinney, J. W. Protein-Protein Interactions in Virus-Host Systems. Front. Microbiol. 2017, 8 (AUG), 1557. https://doi.org/10.3389/fmicb.2017.01557.

(20) Graham, R. L.; Sims, A. C.; Brockway, S. M.; Baric, R. S.; Denison, M. R. The Nsp2 Replicase Proteins of Murine Hepatitis Virus and Severe Acute Respiratory Syndrome Coronavirus Are Dispensable for Viral Replication. J. Virol. 2005, 79 (21), 13399–13411. https://doi.org/10.1128/JVI.79.21.13399-13411.2005.

(21) Angeletti, S.; Benvenuto, D.; Bianchi, M.; Giovanetti, M.; Pascarella, S.; Ciccozzi, M. COVID-2019: The Role of the Nsp2 and Nsp3 in Its Pathogenesis. J. Med. Virol. 2020, 0–3. https://doi.org/10.1002/jmv.25719.

(22) Oostra, M.; te Lintelo, E. G.; Deijs, M.; Verheije, M. H.; Rottier, P. J. M.; de Haan, C. A. M. Localization and Membrane Topology of Coronavirus Nonstructural Protein 4: Involvement of the Early Secretory Pathway in Replication. J. Virol. 2007, 81 (22), 12323–12336. https://doi.org/10.1128/jvi.01506-07.

(23) Gadlage, M. J.; Sparks, J. S.; Beachboard, D. C.; Cox, R. G.; Doyle, J. D.; Stobart, C. C.; Denison, M. R. Murine Hepatitis Virus Nonstructural Protein 4 Regulates Virus-Induced Membrane Modifications and Replication Complex Function. J. Virol. 2010, 84 (1), 280–290. https://doi.org/10.1128/jvi.01772-09.

(24) Oudshoorn, D.; Rijs, K.; Limpens, R. W. A. L.; Groen, K.; Koster, A. J.; Snijder, E. J.; Kikkert, M.; Bárcena, M. Expression and Cleavage of Middle East Respiratory Syndrome Coronavirus Nsp3-4 Polyprotein Induce the Formation of Double-Membrane Vesicles That Mimic Those Associated with Coronaviral RNA Replication. MBio 2017, 8 (6). https://doi.org/10.1128/mBio.01658-17.

(25) Kanjanahaluethai, A.; Chen, Z.; Jukneliene, D.; Baker, S. C. Membrane Topology of Murine Coronavirus Replicase Nonstructural Protein 3. Virology 2007, 361 (2), 391–401. https://doi.org/10.1016/j.virol.2006.12.009.

(26) Christie, D. A.; Lemke, C. D.; Elias, I. M.; Chau, L. A.; Kirchhof, M. G.; Li, B.; Ball, E. H.; Dunn, S. D.; Hatch, G. M.; Madrenas, J. Stomatin-Like Protein 2 Binds Cardiolipin and Regulates Mitochondrial Biogenesis and Function. Mol. Cell. Biol. 2011, 31 (18), 3845–3856. https://doi.org/10.1128/mcb.05393-11.

(27) Panda, D.; Gold, B.; Tartell, M. A.; Rausch, K.; Casas-Tinto, S.; Cherry, S. The Transcription Factor FoxK Participates with Nup98 to Regulate Antiviral Gene Expression. MBio 2015, 6 (2). https://doi.org/10.1128/mBio.02509-14.

(28) Yenamandra, S. P. Expression Profile of Nuclear Receptors Upon Epstein -- Barr Virus Induced B Cell Transformation. Exp. Oncol. 2009, 31 (2), 92–96.

(29) Gavin, A. L.; Huang, D.; Huber, C.; Mårtensson, A.; Tardif, V.; Skog, P. D.; Blane, T. R.; Thinnes, T. C.; Osborn, K.; Chong, H. S.; Kargaran, F.; Kimm, P.; Zeitjian, A.; Sielski, R. L.; Briggs, M.; Schulz, S. R.; Zarpellon, A.; Cravatt, B.; Pang, E. S.; Teijaro, J.; de la Torre, J. C.; O’Keeffe, M.; Hochrein, H.; Damme, M.; Teyton, L.; Lawson, B. R.; Nemazee, D. PLD3 and PLD4 Are Single-Stranded Acid Exonucleases That Regulate Endosomal Nucleic-Acid Sensing. Nat. Immunol. 2018, 19 (9), 942–953. https://doi.org/10.1038/s41590-018-0179-y.

(30) Parks, C. L.; Shenk, T. Activation of the Adenovirus Major Late Promoter by Transcription Factors MAZ and Sp1. J. Virol. 1997, 71 (12), 9600–9607. https://doi.org/10.1128/jvi.71.12.9600-9607.1997.

(31) Ovsyannikova, I. G.; Salk, H. M.; Kennedy, R. B.; Haralambieva, I. H.; Zimmermann, M. T.; Grill, D. E.; Oberg, A. L.; Poland, G. A. Gene Signatures Associated with Adaptive Humoral Immunity Following Seasonal Influenza A/H1N1 Vaccination. Genes Immun. 2016, 17 (7), 371–379. https://doi.org/10.1038/gene.2016.34.

(32) Le, M. X.; Haddad, D.; Ling, A. K.; Li, C.; So, C. C.; Chopra, A.; Hu, R.; Angulo, J. F.; Moffat, J.; Martin, A. Kin17 Facilitates Multiple Double-Strand Break Repair Pathways That Govern B Cell Class Switching. Sci. Rep. 2016, 6. https://doi.org/10.1038/srep37215.

(33) Zhong, B.; Zhang, L.; Lei, C.; Li, Y.; Mao, A. P.; Yang, Y.; Wang, Y. Y.; Zhang, X. L.; Shu, H. B. The Ubiquitin Ligase RNF5 Regulates Antiviral Responses by Mediating Degradation of the Adaptor Protein MITA. Immunity 2009, 30 (3), 397–407. https://doi.org/10.1016/j.immuni.2009.01.008.

(34) Fenech, E. J.; Lari, F.; Charles, P. D.; Fischer, R.; Laétitia-Thézénas, M.; Bagola, K.; Paton, A. W.; Paton, J. C.; Gyrd-Hansen, M.; Kessler, B. M.; Christianson, J. C. Interaction Mapping of Endoplasmic Reticulum Ubiquitin Ligases Identifies Modulators of Innate Immune Signalling. https://doi.org/10.1101/2020.03.18.993998.

(35) Owen, R. P.; Gong, L.; Sagreiya, H.; Klein, T. E.; Altman, R. B. VKORC1 Pharmacogenomics Summary. Pharmacogenetics and Genomics. NIH Public Access October 2010, pp 642–644. https://doi.org/10.1097/FPC.0b013e32833433b6.

(36) Ho, D. V.; Chan, J. Y. Induction of Herpud1 Expression by ER Stress Is Regulated by Nrf1. FEBS Lett. 2015, 589 (5), 615–620. https://doi.org/10.1016/j.febslet.2015.01.026.

(37) Qin, L.; Guo, J.; Zheng, Q.; Zhang, H. BAG2 Structure, Function and Involvement in Disease. Cellular and Molecular Biology Letters. BioMed Central Ltd. September 20, 2016. https://doi.org/10.1186/s11658-016-0020-2.

(38) Chartron, J. W.; Suloway, C. J. M.; Zaslaver, M.; Clemons, W. M. Structural Characterization of the Get4/Get5 Complex and Its Interaction with Get3. Proc. Natl. Acad. Sci. U. S. A. 2010, 107 (27), 12127–12132. https://doi.org/10.1073/pnas.1006036107.

(39) Gottier, P.; Gonzalez-Salgado, A.; Menon, A. K.; Liu, Y. C.; Acosta-Serrano, A.; Bütikofer, P. RFT1 Protein Affects Glycosylphosphatidylinositol (GPI) Anchor Glycosylation. J. Biol. Chem. 2017, 292 (3), 1103–1111. https://doi.org/10.1074/jbc.M116.758367.

(40) Haeuptle, M. A.; Pujol, F. M.; Neupert, C.; Winchester, B.; Kastaniotis, A. J. J.; Aebi, M.; Hennet, T. Human RFT1 Deficiency Leads to a Disorder of N-Linked Glycosylation. Am. J. Hum. Genet. 2008, 82 (3), 600–606. https://doi.org/10.1016/j.ajhg.2007.12.021.

(41) Papp, B.; Brouland, J. P.; Arbabian, A.; Gélébart, P.; Kovács, T.; Bobe, R.; Enouf, J.; Varin-Blank, N.; Apáti, Á. Endoplasmic Reticulum Calcium Pumps and Cancer Cell Differentiation. Biomolecules. MDPI AG 2012, pp 165–186. https://doi.org/10.3390/biom2010165.

(42) Seo, J.; Lee, S. H.; Park, S. Y.; Jeong, M. H.; Lee, S. Y.; Kim, M. J.; Yoo, J. Y.; Jang, S.; Choi, K. C.; Yoon, Ho G. GPR177 Promotes Gastric Cancer Proliferation by Suppressing Endoplasmic Reticulum Stress-Induced Cell Death. J. Cell. Biochem. 2019, 120 (2), 2532–2539. https://doi.org/10.1002/jcb.27545.

(43) Cui, J.; Chen, W.; Sun, J.; Guo, H.; Madley, R.; Xiong, Y.; Pan, X.; Wang, H.; Tai, A. W.; Weiss, M. A.; Arvan, P.; Liu, M. Competitive Inhibition of the Endoplasmic Reticulum Signal Peptidase by Non-Cleavable Mutant Preprotein Cargos. J. Biol. Chem. 2015, 290 (47), 28131–28140. https://doi.org/10.1074/jbc.M115.692350.

(44) Blanc, M.; Hsieh, W. Y.; Robertson, K. A.; Watterson, S.; Shui, G.; Lacaze, P.; Khondoker, M.; Dickinson, P.; Sing, G.; Rodríguez-Martín, S.; Phelan, P.; Forster, T.; Strobl, B.; Müller, M.; Riemersma, R.; Osborne, T.; Wenk, M. R.; Angulo, A.; Ghazal, P. Host Defense against Viral Infection Involves Interferon Mediated Down-Regulation of Sterol Biosynthesis. PLoS Biol. 2011, 9 (3). https://doi.org/10.1371/journal.pbio.1000598.

(45) van ′t Wout, A. B.; Swain, J. V.; Schindler, M.; Rao, U.; Pathmajeyan, M. S.; Mullins, J. I.; Kirchhoff, F. Nef Induces Multiple Genes Involved in Cholesterol Synthesis and Uptake in Human Immunodeficiency Virus Type 1-Infected T Cells. J. Virol. 2005, 79 (15), 10053–10058. https://doi.org/10.1128/jvi.79.15.10053-10058.2005.

(46) Diamond, D. L.; Syder, A. J.; Jacobs, J. M.; Sorensen, C. M.; Walters, K. A.; Proll, S. C.; McDermott, J. E.; Gritsenko, M. A.; Zhang, Q.; Zhao, R.; Metz, T. O.; Camp, D. G.; Waters, K. M.; Smith, R. D.; Rice, C. M.; Katze, M. G. Temporal Proteome and Lipidome Profiles Reveal Hepatitis C Virus-Associated Reprogramming of Hepatocellular Metabolism and Bioenergetics. PLoS Pathog. 2010, 6 (1). https://doi.org/10.1371/journal.ppat.1000719.

(47) Xiao, J.; Li, W.; Zheng, X.; Qi, L.; Wang, H.; Zhang, C.; Wan, X.; Zheng, Y.; Zhong, R.; Zhou, X.; Lu, Y.; Li, Z.; Qiu, Y.; Liu, C.; Zhang, F.; Zhang, Y.; Xu, X.; Yang, Z.; Chen, H.; Zhai, Q.; Wei, B.; Wang, H. Targeting 7-Dehydrocholesterol Reductase Integrates Cholesterol Metabolism and IRF3 Activation to Eliminate Infection. Immunity 2020, 52 (1), 109–122.e6. https://doi.org/10.1016/j.immuni.2019.11.015.

(48) Larance, M.; Kirkwood, K. J.; Xirodimas, D. P.; Lundberg, E.; Uhlen, M.; Lamond, A. I. Characterization of MRFAP1 Turnover and Interactions Downstream of the NEDD8 Pathway. Mol. Cell. Proteomics 2012, 11 (3). https://doi.org/10.1074/mcp.M111.014407.

(49) Kelly, T. J.; Suzuki, H. I.; Zamudio, J. R.; Suzuki, M.; Sharp, P. A. Sequestration of MicroRNA-Mediated Target Repression by the Ago2-Associated RNA-Binding Protein FAM120A. RNA 2019, 25 (10), 1291–1297. https://doi.org/10.1261/rna.071621.119.

(50) Bartolomé, R. A.; García-Palmero, I.; Torres, S.; López-Lucendo, M.; Balyasnikova, I. V.; Casal, J. I. IL13 Receptor Α2 Signaling Requires a Scaffold Protein, FAM120A, to Activate the FAK and PI3K Pathways in Colon Cancer Metastasis. Cancer Res. 2015, 75 (12), 2434–2444. https://doi.org/10.1158/0008-5472.CAN-14-3650.

(51) Lampert, F.; Stafa, D.; Goga, A.; Soste, M. V.; Gilberto, S.; Olieric, N.; Picotti, P.; Stoffel, M.; Peter, M. The Multi-Subunit GID/CTLH E3 Ubiquitin Ligase Promotes Cell Proliferation and Targets the Transcription Factor Hbp1 for Degradation. Elife 2018, 7. https://doi.org/10.7554/eLife.35528.

(52) Cho, K. F.; Branon, T. C.; Rajeev, S.; Svinkina, T.; Udeshi, N. D.; Thoudam, T.; Kwak, C.; Rhee, H.-W.; Lee, I.-K.; Carr, S. A.; Ting, A. Y.; Chan, M.; Biohub, Z.; Francisco, S. Split-TurboID Enables Contact-Dependent Proximity Labeling in Cells. https://doi.org/10.1073/pnas.1919528117/-/DCSupplemental.

(53) Kwak, C.; Shin, S.; Park, J.-S.; Jung, M.; Thi My Nhung, T.; Kang, M.-G.; Lee, C.; Kwon, T.-H.; Ki Park, S.; Young Mun, J.; Kim, J.-S.; Rhee, H.-W. Contact-ID, a Tool for Profiling Organelle Contact Sites, Reveals Regulatory Proteins of Mitochondrial-Associated Membrane Formation. https://doi.org/10.1073/pnas.1916584117/-/DCSupplemental.

(54) Carreras-Sureda, A.; Jaña, F.; Urra, H.; Durand, S.; Mortenson, D. E.; Sagredo, A.; Bustos, G.; Hazari, Y.; Ramos-Fernández, E.; Sassano, M. L.; Pihán, P.; van Vliet, A. R.; González-Quiroz, M.; Torres, A. K.; Tapia-Rojas, C.; Kerkhofs, M.; Vicente, R.; Kaufman, R. J.; Inestrosa, N. C.; Gonzalez-Billault, C.; Wiseman, R. L.; Agostinis, P.; Bultynck, G.; Court, F. A.; Kroemer, G.; Cárdenas, J. C.; Hetz, C. Non-Canonical Function of IRE1α Determines Mitochondria-Associated Endoplasmic Reticulum Composition to Control Calcium Transfer and Bioenergetics. Nat. Cell Biol. 2019, 21 (6), 755–767. https://doi.org/10.1038/s41556-019-0329-y.

(55) Kim, J. H.; Rhee, J. K.; Ahn, D. G.; Kim, K. P.; Oh, J. W. Interaction of Stomatin with Hepatitis C Virus RNA Polymerase Stabilizes the Viral RNA Replicase Complexes on Detergent-Resistant Membranes. J. Microbiol. Biotechnol. 2014, 24 (12), 1744–1754. https://doi.org/10.4014/jmb.1409.09063.

(56) Wintachai, P.; Wikan, N.; Kuadkitkan, A.; Jaimipuk, T.; Ubol, S.; Pulmanausahakul, R.; Auewarakul, P.; Kasinrerk, W.; Weng, W. Y.; Panyasrivanit, M.; Paemanee, A.; Kittisenachai, S.; Roytrakul, S.; Smith, D. R. Identification of Prohibitin as a Chikungunya Virus Receptor Protein. J. Med. Virol. 2012, 84 (11), 1757–1770. https://doi.org/10.1002/jmv.23403.

(57) Too, I. H. K.; Bonne, I.; Tan, E. L.; Chu, J. J. H.; Alonso, S. Prohibitin Plays a Critical Role in Enterovirus 71 Neuropathogenesis. PLoS Pathog. 2018, 14 (1). https://doi.org/10.1371/journal.ppat.1006778.

(58) Richter, K. T.; Kschonsak, Y. T.; Vodicska, B.; Hoffmann, I. FBXO45-MYCBP2 Regulates Mitotic Cell Fate by Targeting FBXW7 for Degradation. Cell Death Differ. 2020, 27 (2), 758–772. https://doi.org/10.1038/s41418-019-0385-7.

(59) Song, X.; Liu, S.; Wang, W.; Ma, Z.; Cao, X.; Jiang, M. E3 Ubiquitin Ligase RNF170 Inhibits Innate Immune Responses by Targeting and Degrading TLR3 in Murine Cells. Cell. Mol. Immunol. 2019. https://doi.org/10.1038/s41423-019-0236-y.

(60) Lu, J. P.; Wang, Y.; Sliter, D. A.; Pearce, M. M. P.; Wojcikiewicz, R. J. H. RNF170 Protein, an Endoplasmic Reticulum Membrane Ubiquitin Ligase, Mediates Inositol 1,4,5-Trisphosphate Receptor Ubiquitination and Degradation. J. Biol. Chem. 2011, 286 (27), 24426–24433. https://doi.org/10.1074/jbc.M111.251983.

(61) Berridge, M. J. Inositol Trisphosphate and Calcium Signalling; 1993.

(62) Bartok, A.; Weaver, D.; Golenár, T.; Nichtova, Z.; Katona, M.; Bánsághi, S.; Alzayady, K. J.; Thomas, V. K.; Ando, H.; Mikoshiba, K.; Joseph, S. K.; Yule, D. I.; Csordás, G.; Hajnóczky, G. IP3 Receptor Isoforms Differently Regulate ER-Mitochondrial Contacts and Local Calcium Transfer. Nat. Commun. 2019, 10 (1), 1–14. https://doi.org/10.1038/s41467-019-11646-3.

(63) Missiroli, S.; Patergnani, S.; Caroccia, N.; Pedriali, G.; Perrone, M.; Previati, M.; Wieckowski, M. R.; Giorgi, C. Mitochondria-Associated Membranes (MAMs) and Inflammation. 2018. https://doi.org/10.1038/s41419-017-0027-2.

(64) Zhou, Y.; Frey, T. K.; Yang, J. J. Viral Calciomics: Interplays between Ca 2+ and Virus. Cell Calcium 2009, 46, 1–17. https://doi.org/10.1016/j.ceca.2009.05.005.

(65) Nieto-Torres, J. L.; Verdiá-Báguena, C.; Jimenez-Guardeño, J. M.; Regla-Nava, J. A.; Castaño-Rodriguez, C.; Fernandez-Delgado, R.; Torres, J.; Aguilella, V. M.; Enjuanes, L. Severe Acute Respiratory Syndrome Coronavirus E Protein Transports Calcium Ions and Activates the NLRP3 Inflammasome. 2015. https://doi.org/10.1016/j.virol.2015.08.010.

(66) Bojkova, D.; Klann, K.; Koch, B.; Widera, M.; Krause, D.; Ciesek, S.; Cinatl, J.; Münch, C. Proteomics of SARS-CoV-2-Infected Host Cells Reveals Therapy Targets. Nature 2020, 1–8. https://doi.org/10.1038/s41586-020-2332-7.

(67) Wright, M. T.; Kouba, L.; Plate, L. Thyroglobulin Interactome Profiling Uncovers Molecular Mechanisms of 1 Thyroid Dyshormonogenesis 2 3. https://doi.org/10.1101/2020.04.08.032482.

(68) Fonslow, B. R.; Niessen, S. M.; Singh, M.; Wong, C. C.; Xu, T.; Carvalho, P. C.; Choi, J.; Park, S. K.; Yates 3rd, J. R. Single-Step Inline Hydroxyapatite Enrichment Facilitates Identification and Quantitation of Phosphopeptides from Mass-Limited Proteomes with MudPIT. J Proteome Res 2012, 11 (5), 2697–2709. https://doi.org/10.1021/pr300200x.

(69) Keilhauer, E. C.; Hein, M. Y.; Mann, M. Accurate Protein Complex Retrieval by Affinity Enrichment Mass Spectrometry (AE-MS) Rather than Affinity Purification Mass Spectrometry (AP-MS). Mol. Cell. Proteomics 2015, 14 (1), 120–135. https://doi.org/10.1074/mcp.M114.041012.

(70) Kuleshov, M. V; Jones, M. R.; Rouillard, A. D.; Fernandez, N. F.; Duan, Q.; Wang, Z.; Koplev, S.; Jenkins, S. L.; Jagodnik, K. M.; Lachmann, A.; Mcdermott, M. G.; Monteiro, C. D.; Gundersen, G. W.; Ma’ayan, A. Enrichr: A Comprehensive Gene Set Enrichment Analysis Web Server 2016 Update. Nucleic Acids Res. 2016, 44. https://doi.org/10.1093/nar/gkw377.

