## Supplemental Information for "Comparative multiplexed interactomics of SARS-CoV-2 and homologous coronavirus non-structural proteins identifies unique and shared host-cell dependencies"

**TABLE OF CONTENT**

Supplementary Figures S1 – S10 pp 3 – 18

Supplementary Tables p. 19

Supplementary References p. 20

### SUPPLEMENTARY FIGURES

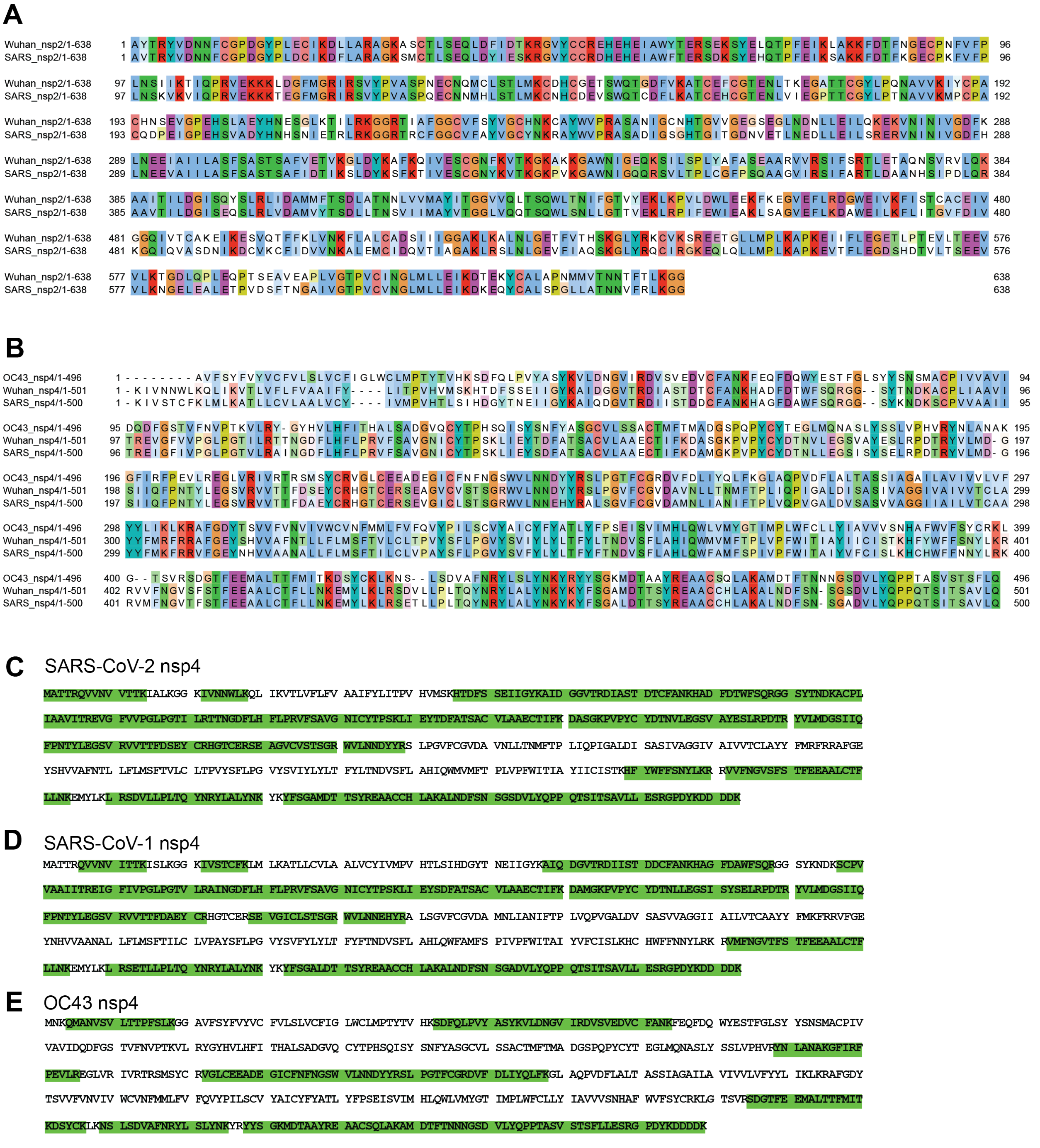


**Figure S1. Amino acid sequence comparison and MS sequence coverage of CoV non-structural protein homologs.**

(A) Amino acid sequence alignment of nsp2 homologs from SARS-CoV-1 and SARS-CoV-2. Identical or similar residues are highlighted and color scheme corresponds to amino acid properties.
(B) Amino acid sequence alignment of nsp4 homologs from SARS-CoV-1, SARS-CoV-2 and OC43. Identical or similar residues are highlighted and color scheme corresponds to amino acid properties.
(C) Tandem MS sequence coverage of SARS-Cov-2 (Wuhan) nsp4. Detected peptides are indicated in green.
(D) Tandem MS sequence coverage of SARS-Cov-1 (Urbani) nsp4. Detected peptides are indicated in green.
(E) Tandem MS sequence coverage of OC43 nsp4. Detected peptides are indicated in green.

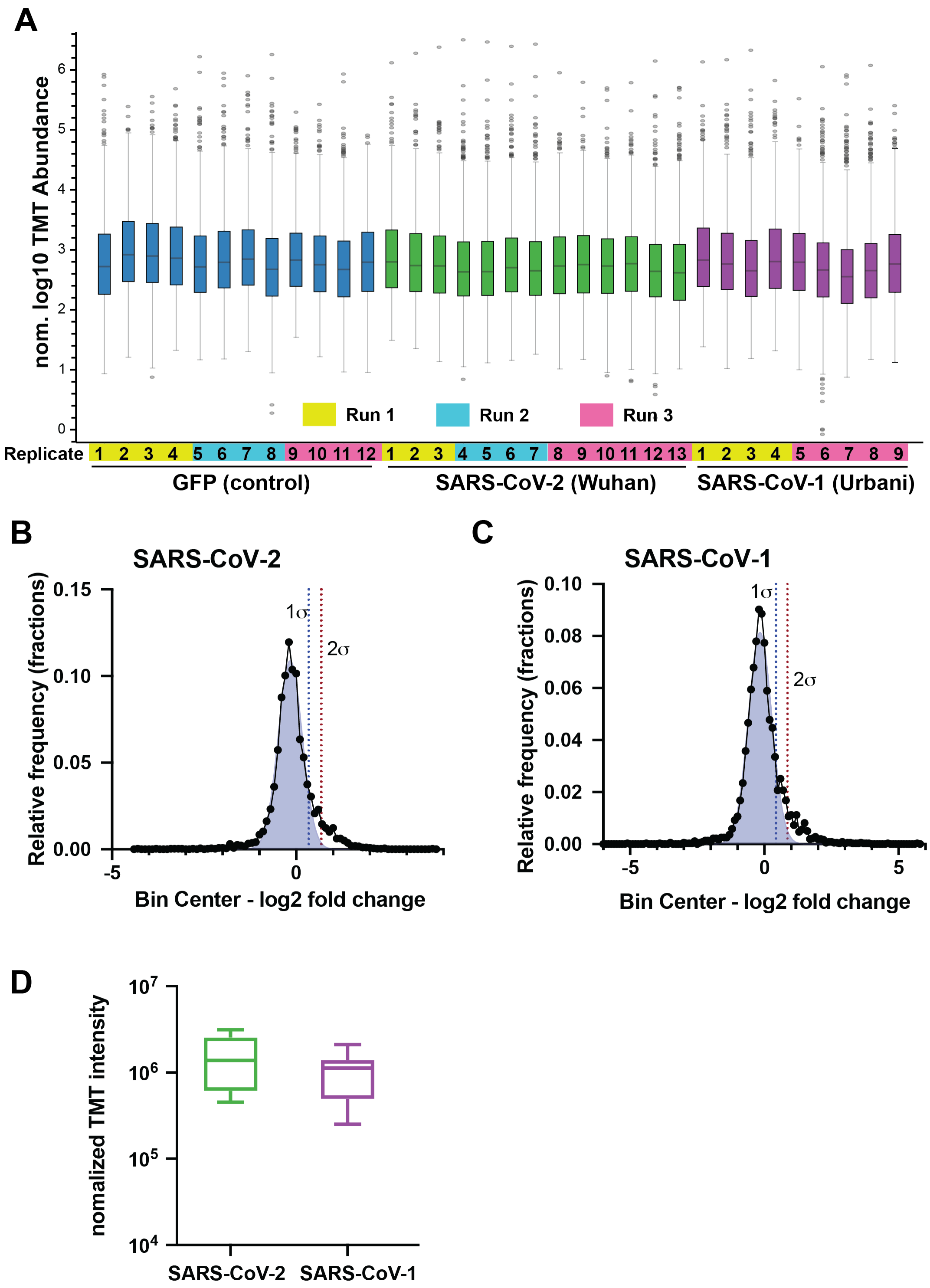


**Figure S2. TMT normalization and filtering of nsp2 interactors.**

(A) Normalized log10 TMT abundances of all proteins in the affinity purification samples expressing GFP control or nsp2 homologs as bait proteins. Pairing of individual samples into three separate mass spectrometry runs is indicated by the color bars.

(B-C) Histogram of log2 fold change enrichment of proteins in nsp2 affinity purification samples compared to GFP controls. A gaussian non-least square fit of the distribution is shown in light blue. The standard deviation of the distribution ($\sigma$) is indicated and 1 $\sigma$ and 2 $\sigma$ (dotted lines) were used for the variable cutoffs to define medium- and high-confidence interactors, respectively.

(B) shows the distribution for SARS-CoV-2 nsp2 with 1 $\sigma$ = 0.5.

(C) shows the distribution for SARS-CoV-2 nsp2 with 1 $\sigma$ = 0.43.

(D) Normalized TMT intensities comparing the abundances of SARS-CoV-2 and SARS-CoV-1 nsp2 homologs in the replicate samples.

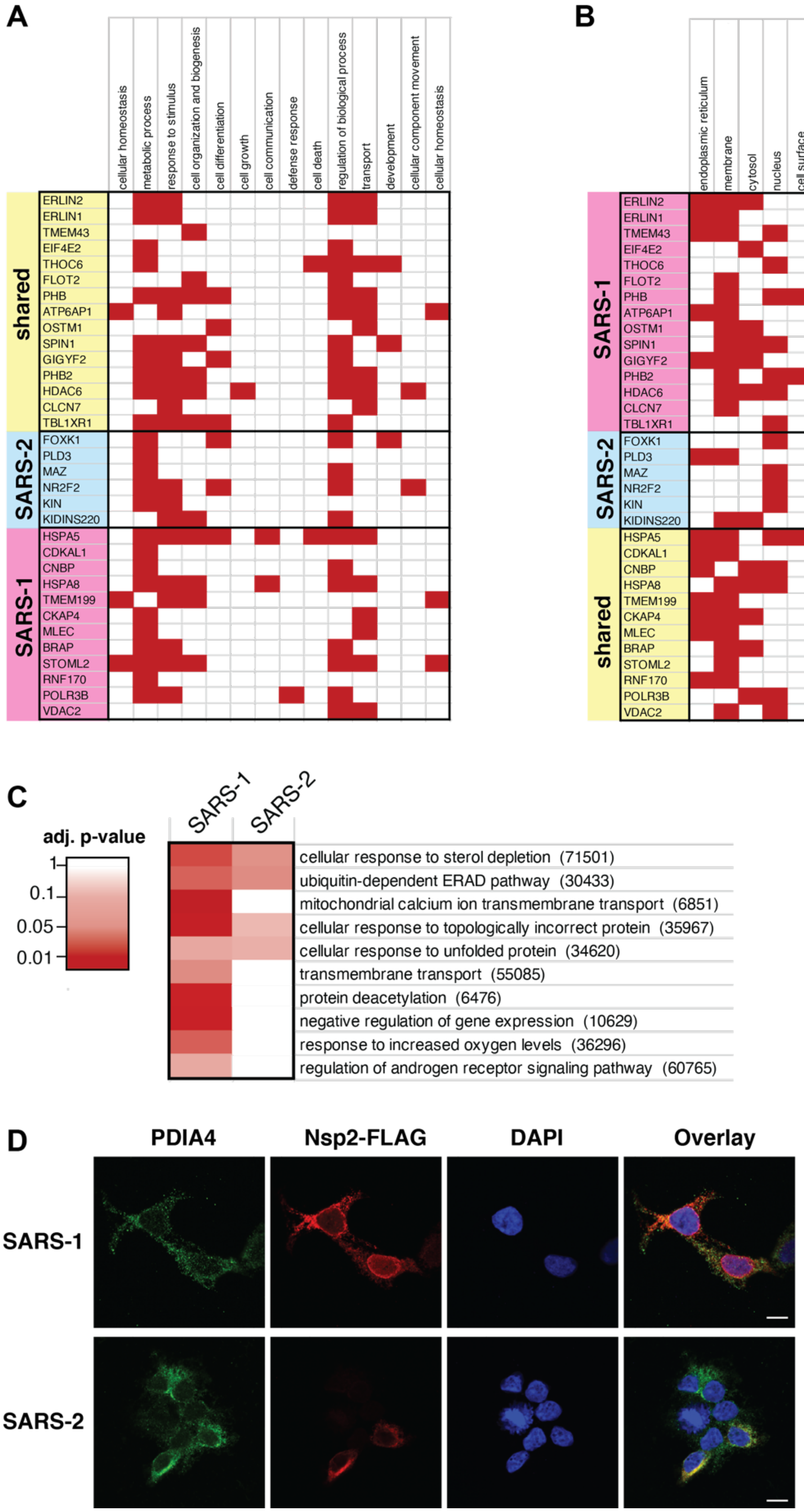


**Figure S3. Gene ontology (GO) pathway analysis of interactors and cellular localization of nsp2 homologs.**

(A-B) GO terms associated with the individual interactors of nsp2. Terms were assigned in the Protein Annotation node in Proteome Discoverer 2.4. Proteins were grouped according to the hierarchical clustering in Fig. 3B to distinguish shared and distinct interactors of SARS-CoV-1 and SARS-CoV-2 nsp2. (A) GO terms for biological processes. (B) GO terms for cellular components.

(C) Comparisons of pathways identified in gene set enrichment analysis of interactors of nsp2 homologs. Gene set enrichment analysis of high-and medium-confidence interactors was performed in EnrichR. GO terms for biological processes with adjusted p-values < 0.1 were included in the analysis and filtered manually to remove redundant terms containing similar genes. Non-redundant pathways are shown and color scheme indicates confidence of enrichment as represented by adjusted p-values of the EnrichR analysis.

(D) Confocal microscopy images of nsp2 homologs expressed in HEK293T cells, stained for PDIA4 (ER marker, green), FLAG-nsp2(red), and DAPI (blue). Scale bar is 10 μm.

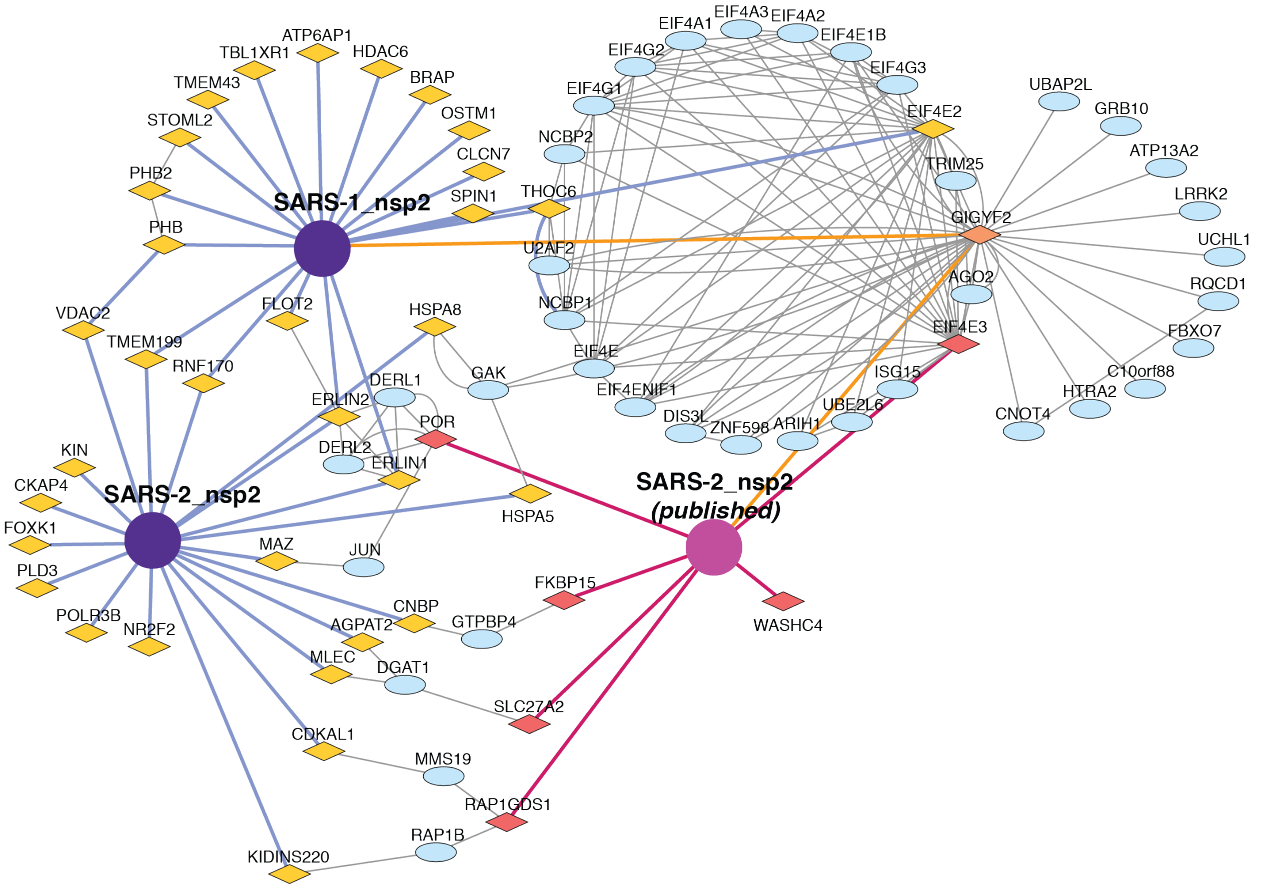


**Figure S4. Nsp2 interactome overlap with published dataset.**
Extended and overlapping interactors between our nsp2 dataset and previously published interactors of SARS-CoV-2 nsp2[1]. Nsp2 proteins are shown as purple circles (our dataset) or pink circle (published dataset). Previously published primary interactors are shown as dark pink diamonds, novel primary interactors identified in this study are shown as yellow diamonds, overlapping primary interactors (GIGYF2) are shown as orange diamonds, and overlapping secondary interactors scraped from the STRING database are shown as blue ellipses. Previously identified primary interactions are shown with red edges, novel primary interactions identified in this study are shown with blue edges, overlapping primary interactors are shown with orange edges, and secondary interactions scraped from the STRING database are shown as grey edges between nodes.

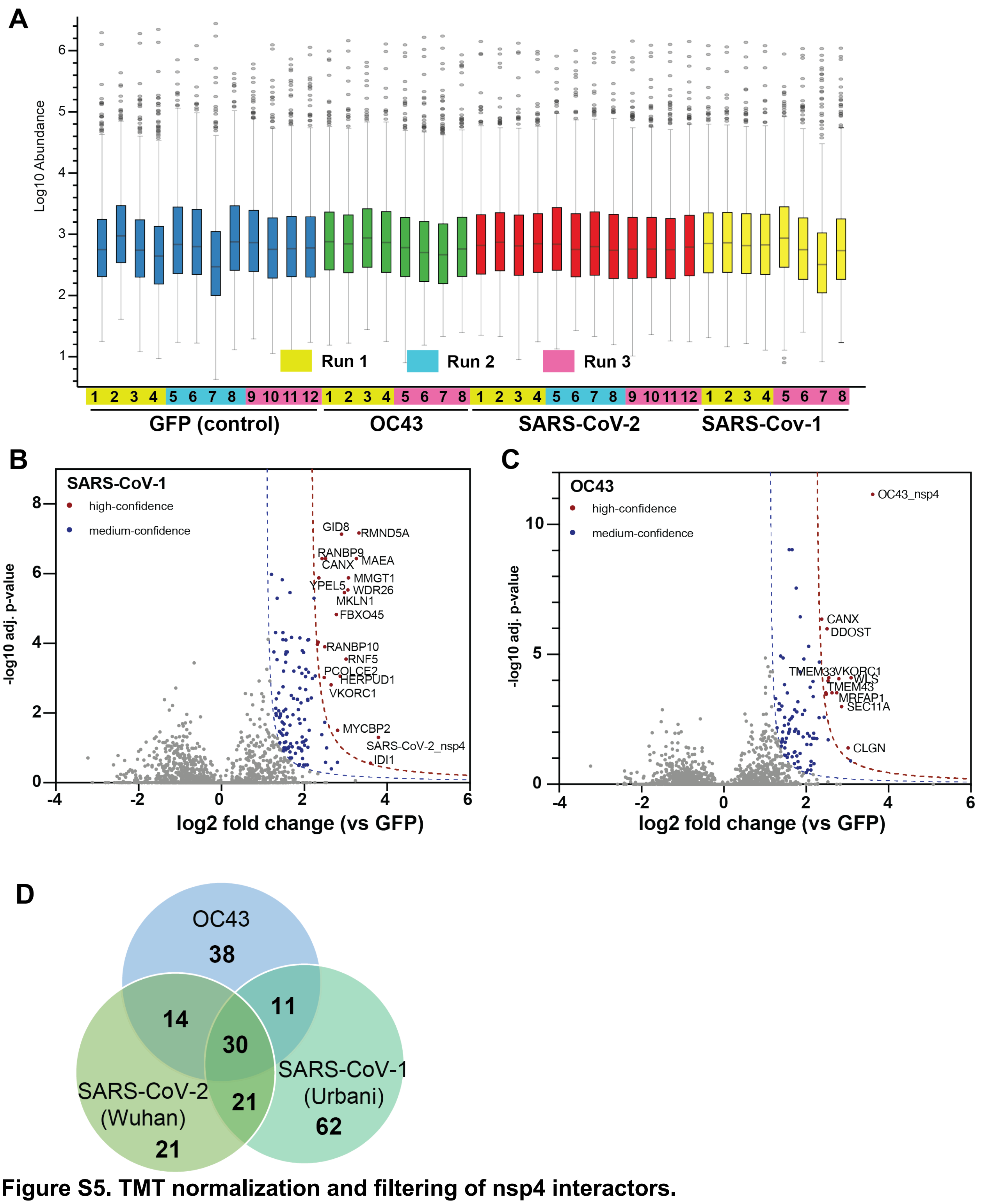


**Figure S5. TMT normalization and filtering of nsp4 interactors.**

(A) Normalized log10 TMT abundances of all proteins in the affinity purification samples expressing GFP control or nsp4 homologs as bait proteins. Pairing of individual samples into three separate mass spectrometry runs is indicated by the color bars.

(B-C) Volcano plot of SARS-CoV-1 (B) and OC43 nsp4 (C) interactors to identify medium- and high-confidence interactors. Plotted are log2 TMT intensity fold changes for proteins between nsp2 bait channels and GFP mock transfections versus -log10 adjusted p-values. Curves for the variable cutoffs used to define high-confidence (red) or medium confidence (blue) interactors are shown. 1$\sigma$ = 0.4 for (B), 1 $\sigma$ = 0.40 for (C).

(C) Venn diagram comparing medium-confidence interactors of nsp4 homologs.

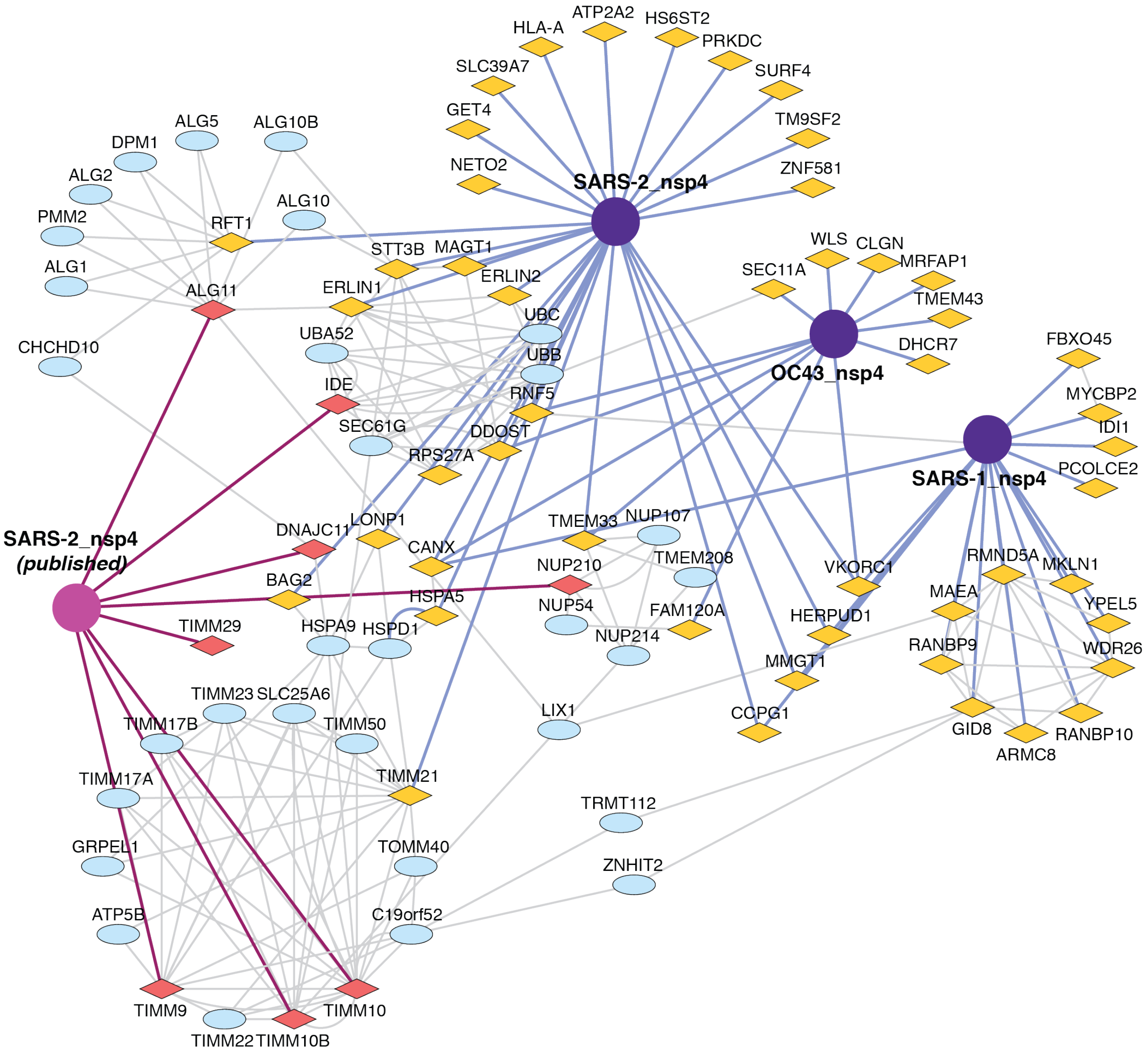


**Figure S6. Nsp2 interactome overlap with published dataset.**
Extended and overlapping interactors between our nsp4 dataset and previously published interactors of SARS-CoV-2 nsp4[1]. Nsp4 proteins are shown as purple circles (our dataset) or pink circle (published dataset). Previously published primary interactors are shown as dark pink diamonds, novel primary interactors identified in this study are shown as yellow diamonds, overlapping primary interactors are shown as orange diamonds, and overlapping secondary interactors scraped from the STRING database are shown as blue ellipses. Previously identified primary interactions are shown with red edges, novel primary interactions identified in this study are shown with blue edges, and secondary interactions scraped from the STRING database are shown as grey edges between nodes. The extended overlapping interactome reveals each Nsp4 protein uniquely plugs into clusters of proteins involved in the same pathway, which appear as circles of nodes such as the TIMM pathway involved in protein import into the mitochondria on the bottom left.

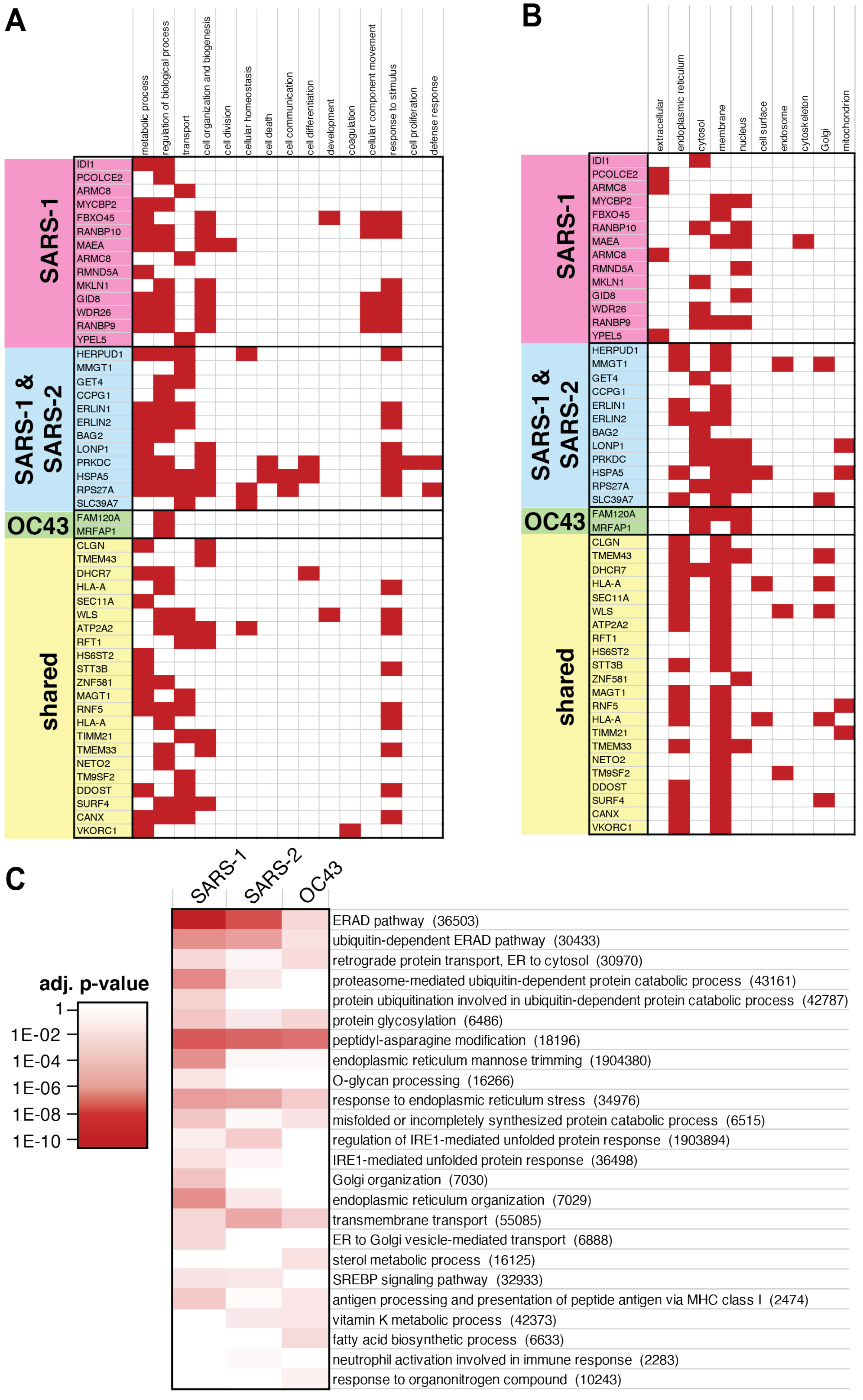


**Figure S7. Gene ontology (GO) pathway analysis of nsp4 homolog interactors.**

(A-B) GO terms associated with the individual interactors of nsp4. Terms were assigned in the Protein Annotation node in Proteome Discoverer 2.4. Proteins were grouped according to the hierarchical clustering in Fig. 4C to distinguish shared and distinct interactors of SARS-CoV-1, SARS-CoV-2, and OC43 nsp4. (A) GO terms for biological processes. (B) GO terms for cellular components.

(C) Comparisons of pathways identified in gene set enrichment analysis of interactors of nsp4 homologs. Gene set enrichment analysis of high-and medium-confidence interactors was performed in EnrichR. GO terms for biological processes with adjusted p-values < 0.1 were included in the analysis and filtered manually to remove redundant terms containing similar genes. Non-redundant pathways are shown and color scheme indicates confidence of enrichment as represented by adjusted p-values of the EnrichR analysis.

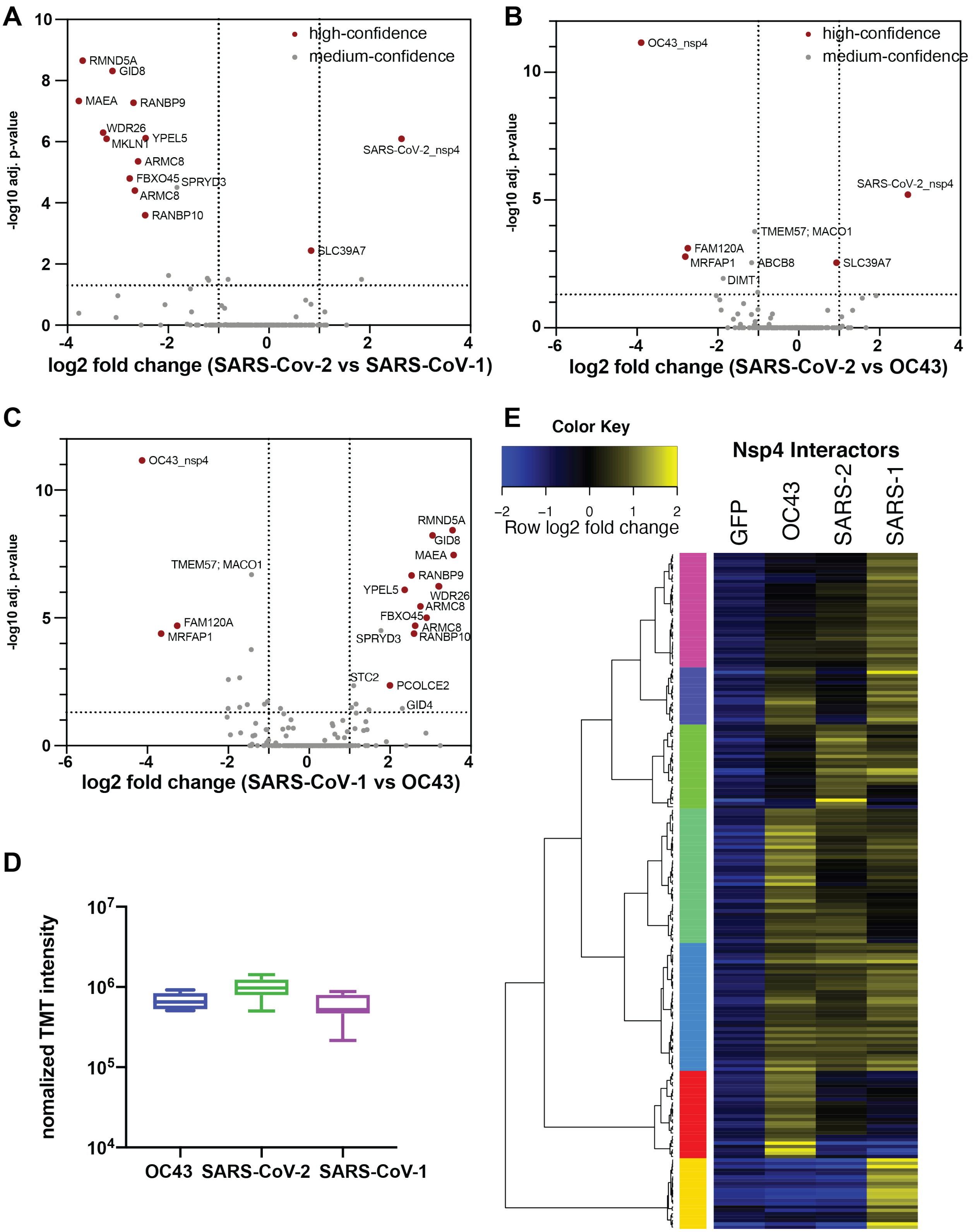


**Figure S8. Comparative analysis of nsp4 homologs.**

(A-C) Volcano plot comparing interactions between nsp4 homolog from SARS-CoV-1, SARS-CoV-2, and OC43. Only high- and medium confidence interactors of nsp4 are shown and high confidence interactors are highlighted in red. (A) Comparison of SARS-CoV-1 and SARS-CoV-2. (B) Comparison of SARS-CoV-2 and OC43. (C) Comparison of SARS-CoV-1 and OC43.

(D) Normalized TMT intensities comparing the abundances of SARS-CoV-2, SARS-CoV-1, and OC43 nsp4 homologs in the replicate affinity purification samples.

(E) Heatmap of high- and medium-confidence nsp4 homolog interactors compared to GFP control. log2 fold change is color-coded and centered by row (blue low, yellow high enrichment). Hierarchical clustering using Ward's method shown on the left was carried out on euclidean distances of log2 fold changes scaled by row.

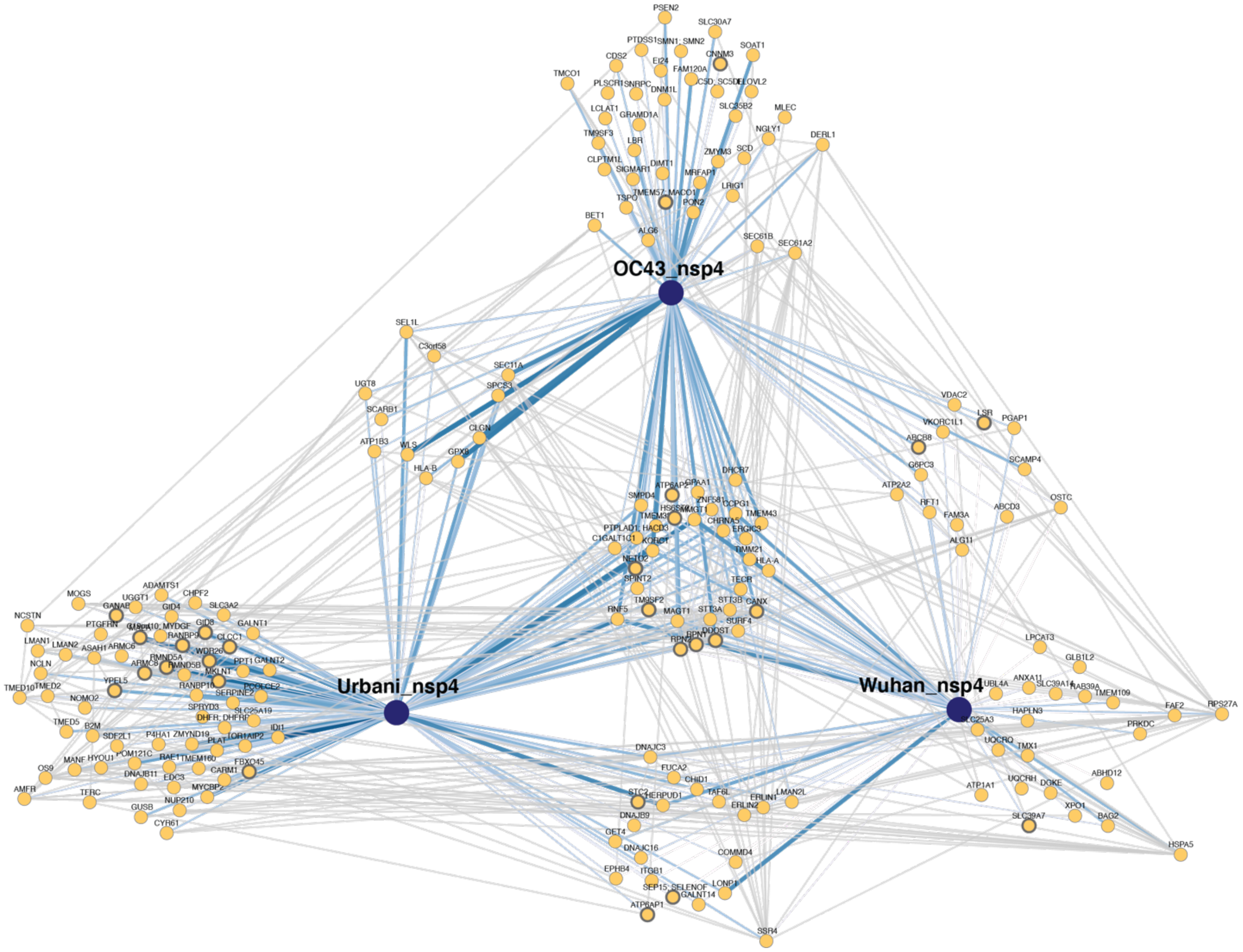


**Figure S9. Network map of high- and medium-confidence nsp4 homolog interactors.**

PPI network map of high- and medium-confidence interactors of nsp4 homolog. Blue lines indicate viral-host PPIs, where line width corresponds to fold enrichment compared to the GFP control. Grey lines indicate annotated host-host PPIs in STRING (score > 0.75).

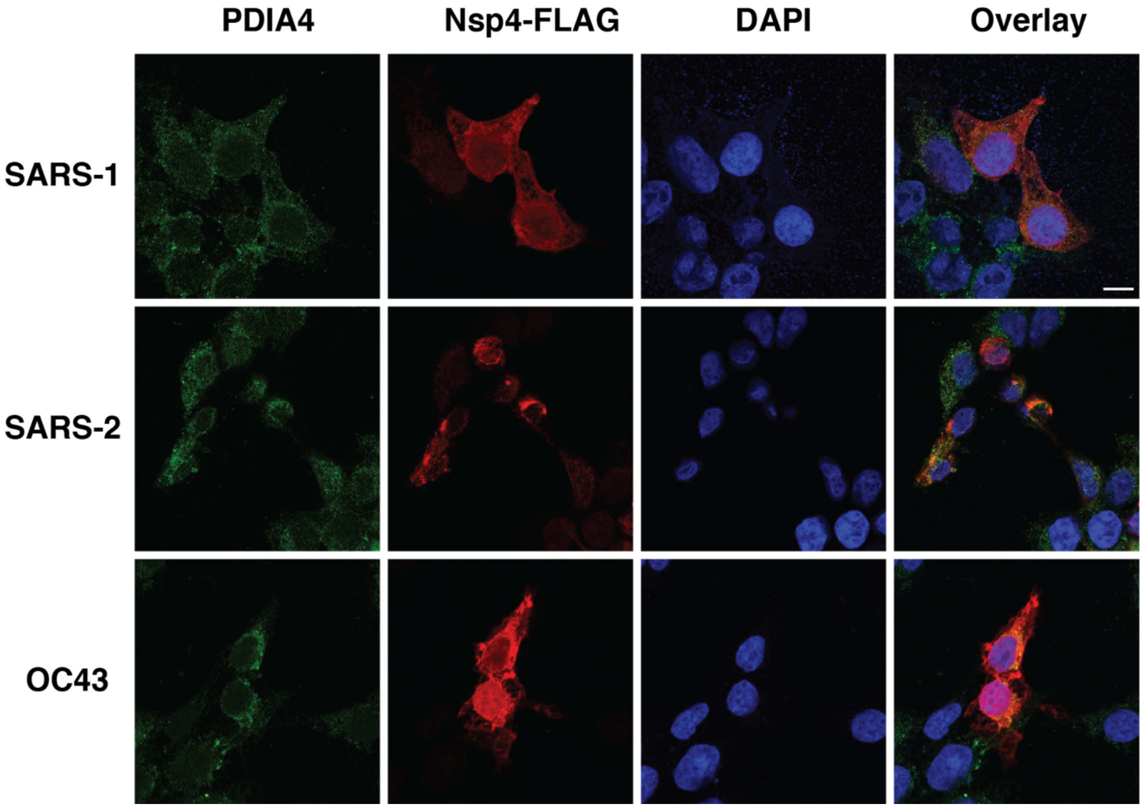


**Figure S10. Subcellular localization of nsp4 homologs.**

Confocal immunofluorescence microscopy images of nsp4 homologs expressed in HEK293T cells, stained for PDIA4 (ER marker, green), FLAG-tag (red), and DAPI (blue). Scale bar is 10 µm.

### SUPPLEMENTARY TABLES

(all supplementary tables are available as separate Excel files)

**Table S1.** Proteomics data of comparative nsp2 interactome profiling. Included are protein identifications, quantifications, abundance ratios, statistical analysis, and filtering of medium- and high confidence interactors.

**Table S2.** Comparison of protein abundances between SARS-CoV-2 and SARS-CoV-1 nsp2 interactors.

**Table S3**. Proteomics data of comparative nsp4 interactome profiling. Included are protein identifications, quantifications, abundance ratios, statistical analysis, and filtering of medium- and high confidence interactors.

**Table S4.** Comparison of protein abundances between SARS-CoV-2, SARS-CoV-1, and OC43 nsp2 interactors.

**Table S5.** List of peptide identifications, quantifications, and protein mapping for comparative nsp2 interactome profiling.

**Table S6.** List of peptide identifications, quantifications, and protein mapping for comparative nsp4 interactome profiling.

### SUPPLEMENTARY REFERENCES

(1) Gordon, D. E.; Jang, G. M.; Bouhaddou, M.; Xu, J.; Obernier, K.; White, K. M.; O'Meara, M. J.; Rezelj, V. V.; Guo, J. Z.; Swaney, D. L.; Tummino, T. A.; Huettenhain, R.; Kaake, R. M.; Richards, A. L.; Tutuncuoglu, B.; Foussard, H.; Batra, J.; Haas, K.; Modak, M.; Kim, M.; Haas, P.; Polacco, B. J.; Braberg, H.; Fabius, J. M.; Eckhardt, M.; Soucheray, M.; Bennett, M. J.; Cakir, M.; McGregor, M. J.; Li, Q.; Meyer, B.; Roesch, F.; Vallet, T.; Mac Kain, A.; Miorin, L.; Moreno, E.; Naing, Z. Z. C.; Zhou, Y.; Peng, S.; Shi, Y.; Zhang, Z.; Shen, W.; Kirby, I. T.; Melnyk, J. E.; Chorba, J. S.; Lou, K.; Dai, S. A.; Barrio-Hernandez, I.; Memon, D.; Hernandez-Armenta, C.; Lyu, J.; Mathy, C. J. P.; Perica, T.; Pilla, K. B.; Ganesan, S. J.; Saltzberg, D. J.; Rakesh, R.; Liu, X.; Rosenthal, S. B.; Calviello, L.; Venkataramanan, S.; Liboy-Lugo, J.; Lin, Y.; Huang, X. P.; Liu, Y. F.; Wankowicz, S. A.; Bohn, M.; Safari, M.; Ugur, F. S.; Koh, C.; Savar, N. S.; Tran, Q. D.; Shengjuler, D.; Fletcher, S. J.; O'Neal, M. C.; Cai, Y.; Chang, J. C. J.; Broadhurst, D. J.; Klippsten, S.; Sharp, P. P.; Wenzell, N. A.; Kuzuoglu, D.; Wang, H. Y.; Trenker, R.; Young, J. M.; Cavero, D. A.; Hiatt, J.; Roth, T. L.; Rathore, U.; Subramanian, A.; Noack, J.; Hubert, M.; Stroud, R. M.; Frankel, A. D.; Rosenberg, O. S.; Verba, K. A.; Agard, D. A.; Ott, M.; Emerman, M.; Jura, N.; von Zastrow, M.; Verdin, E.; Ashworth, A.; Schwartz, O.; D'Enfert, C.; Mukherjee, S.; Jacobson, M.; Malik, H. S.; Fujimori, D. G.; Ideker, T.; Craik, C. S.; Floor, S. N.; Fraser, J. S.; Gross, J. D.; Sali, A.; Roth, B. L.; Ruggero, D.; Taunton, J.; Kortemme, T.; Beltrao, P.; Vignuzzi, M.; García-Sastre, A.; Shokat, K. M.; Shoichet, B. K.; Krogan, N. J. A SARS-CoV-2 Protein Interaction Map Reveals Targets for Drug Repurposing. *Nature* **2020**. https://doi.org/10.1038/s41586-020-2286-9.
